# Multilayered molecular and cellular mechanisms underlying intestinal epithelial regeneration

**DOI:** 10.1101/2025.07.20.665802

**Authors:** Dong Woo Seo, Seungsoo Kim, Hoibin Jeong, Mi Hyeon Cho, Ming Gyeong Lee, Jea Hwa Jang, Jun-Seok Lee, Sang-Hyun Choi, Dong-Hoon Kim, Jungmin Choi, Yong Taek Jeong

## Abstract

Intestinal epithelial cells (IECs) are replenished by intestinal stem cells (ISCs) residing in the intestinal crypts and isthmus. Under homeostatic conditions, Lgr5^+^ crypt base columnar cells (CBCs) are considered the primary ISCs^1,2^. The supply of IECs, however, remains continuous even in the absence of Lgr5^+^ CBCs^3^, implying the existence of alternative ISC populations. Over the past decade, two contrasting models have emerged to explain the identities of these compensatory cells. The distinct cell population model proposes that slow-cycling, quiescent cells near the +4 position, act as reserve stem cells (rISCs)^3-10^. The cellular plasticity model suggests a broad spectrum of IECs can dedifferentiate into intermediate fetal-like states, thereby acquiring stemness^11,12^. It is unclear which of these models is correct because the signaling mechanisms governing crypt regenerative responses remain poorly understood. Here, we demonstrate that Hedgehog (Hh) signaling orchestrates intestinal regeneration following Lgr5^+^ CBC loss by uniquely driving the emergence of de novo Lgr5^+^ CBCs from non-Lgr5^+^ CBCs clones. This regenerative mechanism is fundamentally distinct from both classical self-renewal of existing Lgr5^+^ CBCs and fetal-like dedifferentiation in intestinal villi. By accelerating the cell cycle and driving clonal competition among stem cell pools, Hh signaling leaves rare dormant stem cells in the crypt and isthmus regions. These rare cells act as damage-resistant “seeds” that can promote damage recovery, thereby conferring damage resistance on the tissue as a whole. Collectively, our findings identify novel molecular and cellular mechanisms underlying ISC regeneration and suggest potential therapeutic strategies for various intestinal epithelial diseases.

## Introduction

Intestinal epithelial cells (IECs) undergo rapid and continuous replacement^13^. Early experiments found that this cell proliferation was restricted to specific regions of the intestinal crypts, implying the presence of specialized cells responsible for generating IECs^14,15^. These early insights have since been verified with modern molecular genetic techniques, including tamoxifen-inducible Cre-lox systems. The identification and lineage tracing of the Lgr5^+^ crypt base columnar cells (CBCs) was a particularly significant discovery because these multipotent cells possess self-renewal and can differentiate into most IEC types^1^. Moreover, the discovery that Lgr5^+^ CBCs can be cultured in vitro when provided with the appropriate stem cell niche established them as bona fide intestinal stem cells (ISCs)^2^. Nevertheless, genetic ablation of Lgr5^+^ CBCs with diphtheria toxin failed to impair IEC replenishment or epithelial integrity, implying the existence of an alternative cell population capable of compensating for the loss of Lgr5^+^ CBCs^3^.

Inspired by the successful lineage tracing of Lgr5^+^ CBCs, other researchers performed similar experiments searching for the cells responsible for replenishing IECs in the absence of Lgr5^+^ CBCs^3^. Although many genes have been proposed as specific markers for these cells, there is still no consensus on their cellular identity. There are currently two opposing models at the center of an ongoing debate. The reserve stem cell model posits a population of slow-cycling, quiescent stem cells residing above the crypts that compensate for Lgr5^+^ CBC loss. Lineage tracing studies were used to confirm that rare populations of cells expressing markers such as Bmi1^3,4^, Lrig^5^, Tert^6,10^, or Hopx^7^ can give rise to clonal labeling following Lgr5^+^ CBC injury. Recent evidence has raised questions, however, not only about the specificity of these markers, but even about the existence of a genuine reserve stem cell population. Indeed, Lgr5^+^ CBCs loss can lead to clonal traces arising from other typical IEC types, such as early progenitors committed to the mature absorptive^16,17^ or secretory cell lineages^17,18^, as well as terminally differentiated enteroendocrine cells (EECs)^19^ or Paneth cells (PCs)^20^. These findings suggested the cellular plasticity-related dedifferentiation model and fueled an intense debate over the true cellular identities responsible for IEC maintenance upon loss of Lgr5^+^ CBCs.

Regardless of the exact mechanism, not only are differentiated IECs continuously maintained, but the lost Lgr5^+^ CBCs eventually regenerate as well. These regenerated Lgr5^+^ CBCs are crucial for reconstructing the entire crypt-villus unit structure with all its constituent cells^21,22^. There is, however, still a piece missing in our understanding of how Lgr5^+^ CBCs regenerate. In contrast to the well-established homeostatic stem cell niches supporting Lgr5^+^ CBC self-renewal that require activation of various signaling pathways (i.e., Wnt, BMP, Notch, and EGF^23,24^), the niches for Lgr5^+^ CBC regeneration remain largely unexplored.

In this study, therefore, we aimed to identify the specific signaling pathways that mediate recovery from Lgr5^+^ CBC loss and determine how they contribute to the overall regenerative response. We focused primarily on Hedgehog (Hh) signaling because, although it plays a critical role in intestinal development^25-29^, it has not been extensively studied in adult ISCs. Hh signaling has been implicated in regenerative processes in other organs, including brain^24,30^, hair follicle^31^, and colon^32^. In addition, although they did not clarify the details of the underlying mechanism, one group recently found that an Hh activator was capable of conferring damage resistance to intestinal organoids^33^.

## Results

### Hh signaling is required for the regenerative response following loss of Lgr5^+^ CBCs

We first investigated whether Hh signaling is required for the re-emergence of Lgr5^+^ CBCs following their depletion. Using *Lgr5-DTA* mice, in which tamoxifen injection induces the expression of diphtheria toxin A (DTA) in *Lgr5*-expressing cells, we successfully induced transient depletion of Lgr5^+^ CBCs by recovery day 1 (rec 1) (Fig. 1a-c). Consistent with previous reports^6,10,34^, we observed robust expression of Tert and Ki-67 in the crypts as part of the regenerative response at rec 1, followed by the re-emergence of Lgr5^+^ CBCs by rec 7 (Fig. 1c,d). When we co-injected tamoxifen with the Hh inhibitor cyclopamine, we did not observe Tert or Ki-67 expression on rec 1, nor did we observe the re-emergence of Lgr5^+^ ISCs by rec 7 (Fig. 1c,d). Mice injected with cyclopamine also showed cleaved-Caspase 3 responses, reduced intestine length, and reduced body weight compared to the genetic and tamoxifen-treated controls at rec 1 (Extended Data Fig. 1).

**Fig. 1.**
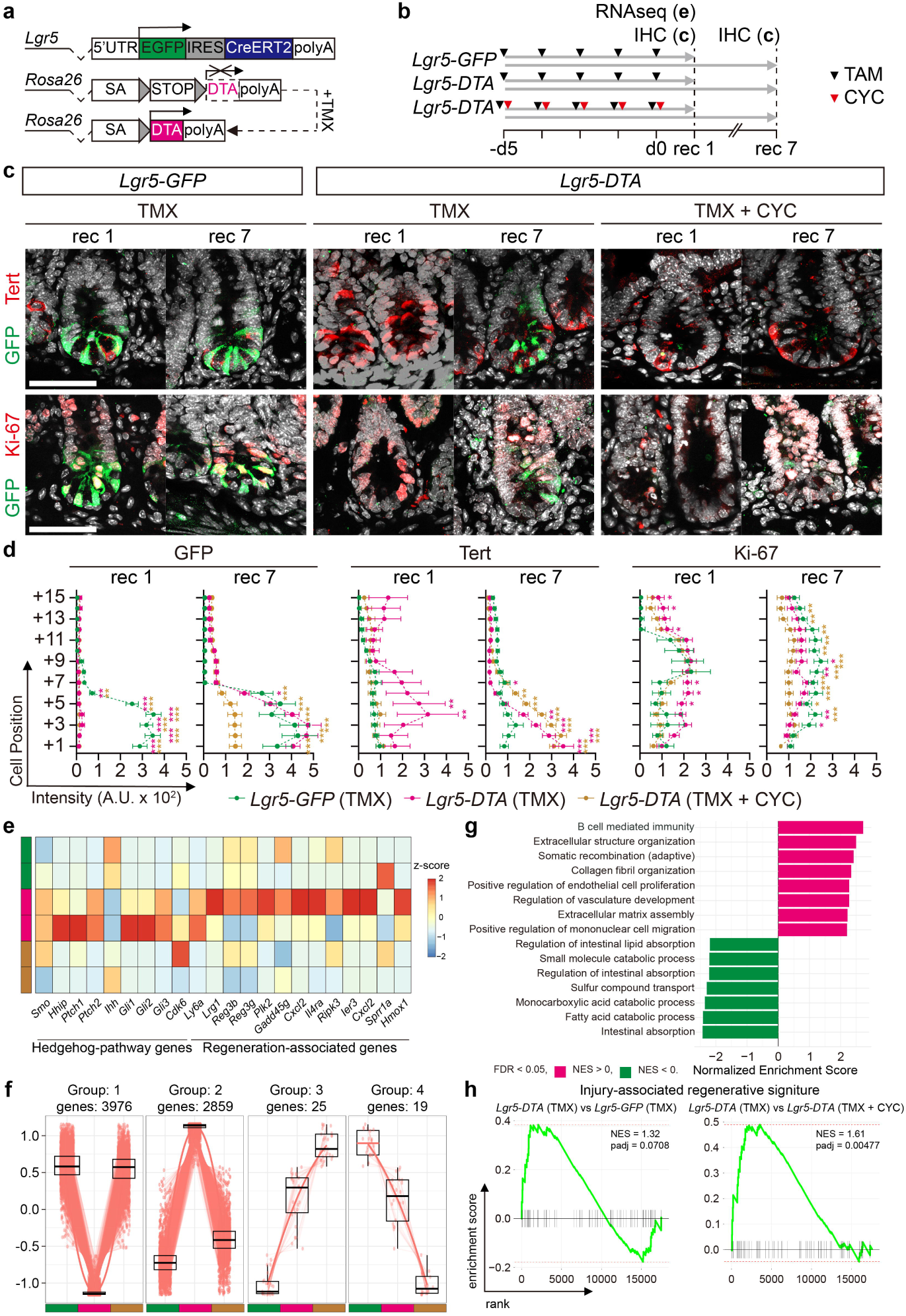
Hh signaling is necessary for the regenerative response triggered by Lgr5^+^ CBC loss. **a**, Schematic showing the genetic constructs for Lgr5^+^ CBC ablation. **b**, Experimental timeline. TMX, tamoxifen; CYC, cyclopamine; rec, recovery days. **c**, Representative images of intestinal crypts showing GFP, Tert, and Ki-67 expression in *Lgr5-GFP*, *Lgr5-DTA*, and *Lgr5-DTA* with CYC treatment at rec 1 and rec 7. *n* = 5 mice per group. Scale bars: 50 µm. **d**, Quantification of GFP, Tert, and Ki-67 immunofluorescence intensity across cell positions at rec 1 and rec 7. *Lgr5-GFP* (green), *Lgr5-DTA* (yellow), *Lgr5-DTA*+CYC (pink). GFP (rec 1, *n* = 36, 40, 25; rec 7, *n* = 29, 29, 36), Tert (rec 1, *n* = 38, 35, 25; rec 7, *n* = 22, 29, 35), Ki-67 (rec 1, *n* = 35, 39, 33; rec 7, *n* = 38, 32, 39). Data are presented as means□±□S.E.M. n = 5 mice per group. **e**–**h**, Bulk RNA-seq analysis of intestinal crypts at rec 1 (*n* = 2 per group). **e**, Heatmap showing normalized expression z-scores of Hh-pathway genes and regeneration-associated genes across groups. **f**, Statistically significant GSEA results of upregulated (pink) or downregulated (green) gene sets from GOBP in Lgr5-DTA (TMX) crypts compared to Lgr5-GFP (TMX). **g**, z-score profiles of differentially expressed genes (DEGs) identified by the likelihood ratio test (LRT) clustered by similar expression dynamics across groups. **h**, GSEA showing enrichment of injury-associated regenerative signatures in Lgr5-DTA (TMX) relative to other groups. Two-way ANOVA with post hoc Tukey’s multiple comparisons test (d). \**P* < 0.05, \*\**P* < 0.01, and \*\*\**P* < 0.001.

In a bulk RNA-seq analysis of isolated crypt epithelium, we found that transient loss of Lgr5^+^ CBCs induced the Hh pathway-related transcripts *Gli1*, *Gli2*, and *Hhip* and broadly altered transcriptomic landscape (Fig. 1e, Extended Data Fig. 2a, b). Gene Set Enrichment Analysis (GSEA) revealed a significant downregulation of genes related to absorption, transport, and digestion, and upregulation of genes related to immune response and tissue remodeling (Fig. 1f, Extended Data Fig. 2c). Notably, these transcriptomic changes were largely abolished by cyclopamine treatment (Fig. 1g). GSEA further revealed that cyclopamine treatment abolished the induction of an injury-associated regenerative signature following transient loss of Lgr5^+^ CBCs. (Fig. 1h). These data indicate that Hh signaling is both activated by transient depletion of Lgr5^+^ CBCs and required for the regeneration that occurs in response to it.

### Activation of Hh signaling expands the pool of Lgr5^+^ CBCs

We next examined the effects of Hh signaling activation on intestinal crypts. To pharmacologically activate Hh signaling, we injected *Lgr5-eGFP-IRES-CreERT2* mice with Smoothened agonist (SAG) (Fig. 2a). This caused a rapid and dynamic remodeling of the crypt epithelium (Fig. 2b-d). On dpi 1, we observed a reduction in Lgr5^+^ CBCs, with those remaining rapidly migrating to the bottom of crypt. On dpi 2, the remaining *Lgr5^+^* CBCs migrated upward, returning to their original location. Finally, on dpi 3, we observed even more Lgr5^+^ CBCs with stronger GFP reporter intensity in a broader distribution than in the homeostatic condition (Fig 2b, c). We also found that Tert expression increased gradually until dpi 2, but then rapidly disappeared at dpi 3. Ki-67 expression, in contrast, continued to increase until dpi 3. Consistent with our histologic results, we found using qRT-PCR that SAG injection increased the levels of the Hh targets *Gli1* and *Gli2*, the ISC markers *Lgr5* and *Olfm4*, and the regenerative marker *Tert* (Extended Data Fig. 3a). Notably, we found SAG injection also increased crypt fission (Fig. 2d), which occurs only rarely in the adult intestine^35^.

**Fig. 2.**
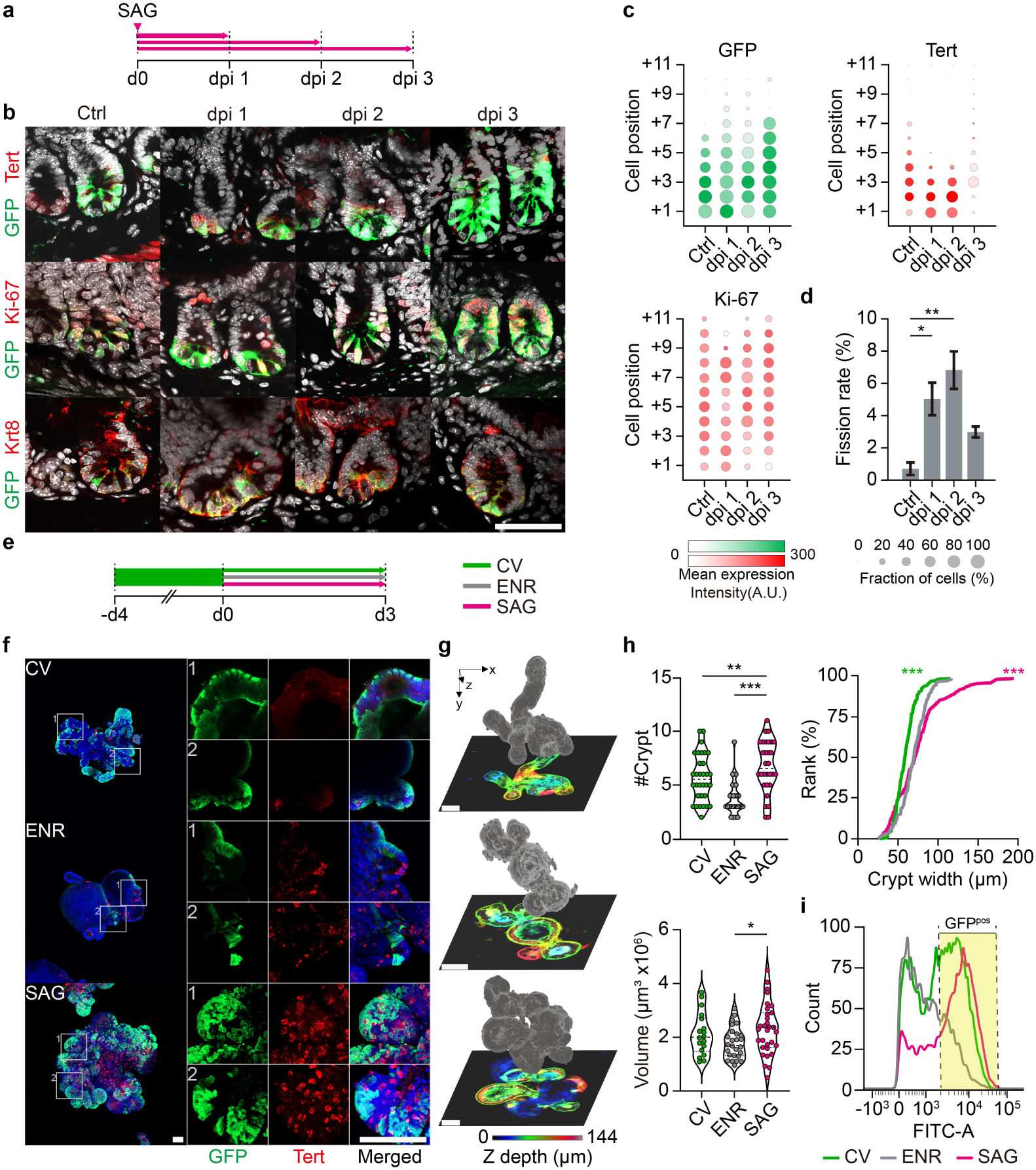
SAG expands the intestinal stem cell pool. **a**, Experimental timeline. SAG, Smoothened agonist; Sac, sacrifice; dpi, days post-injection. **b**, Representative images of intestinal crypts showing GFP, Tert, Ki-67, and Krt8 expression in control and SAG-treated mice at dpi 1, 2, and 3. *n* = 5 mice per group. Scale bars, 50 µm. **c**, Dot plots showing anatomic position and expression intensity of GFP, Tert, and Ki-67 at different timepoints. (GFP, *n* = 35, 30, 37, 34; Tert, *n* = 35, 31, 37, 35; Ki-67, *n* = 31, 32, 29, 33). **d**, Crypt fission rates (*n* = 364, 417, 368, 224). Data are presented as means ± S.E.M. **e**, Organoid culture timeline. ENR, basal culture conditions; CV, Lgr5^+^ CBC enriching conditions; SAG, Hh pathway activation. **f**, Representative images showing GFP and Tert expression in CV, ENR, and SAG-treated organoids. Scale bars: 50 µm. Insets 1 and 2 are magnified and channel-split on the right. **g**, 3D-reconstructed holotomographic images of organoids with depth color-coded from the apical (blue) to basal (red) side. Scale bars, 50 µm. **h**, Quantification of crypts per organoid (*n* = 33, 30, 32), organoid volume (*n* = 19, 27, 31), and organoid crypt width (*n* = 224, 224, 224). **i**, Flow cytometry quantification of GFP^+^ cells. All organoid experiments were performed at least in triplicate. \**P* < 0.05, \*\**P* < 0.01, and \*\*\**P* < 0.001. Ordinary one-way analysis of variance (ANOVA) followed by Tukey’s (**d**) or Dunnett’s (**h**; crypts per organoid, volume) post hoc test. An unpaired t-test with Kolmogorov-Smirnov (**h**; crypt width). \**P* < 0.05, \*\**P* < 0.01, and \*\*\**P* < 0.001.

Next, we used an organoid assay to determine whether Hh signaling directly influences IECs (Fig. 2e). Like CV-treated organoids, in which Lgr5^+^ CBCs undergo self-renewal, SAG-treated organoids exhibited a significant crypt region expansion (Fig. 2f-h) with increased Lgr5^+^ CBCs compared to ENR-treated organoids (Fig 2i). The morphology of these SAG-treated organoids, however, differed markedly from that of CV-treated organoids. While CV-treated organoids showed narrower, elongated crypts, SAG-treated organoids produced broader crypt regions accompanied by an overall increase in organoid volume (Fig 2h). Using high-resolution holotomography, we compared the real-time morphological changes in CV-, ENR-, and SAG-treated organoids (Fig. 2g, Extended Data Fig. 3b, Supplementary Video 1). While CV-treated organoids exhibited rapid crypt apex elongation compared to ENR-treated organoids, SAG-treated organoids broadened at the isthmus region and transiently induced satellite crypts from there. Collectively, these data indicate Hh signaling is sufficient to directly drive dynamic changes in Lgr5^+^ CBCs via mechanisms distinct from their self-renewal.

### Activation of Hh signaling facilitates Lgr5^+^ CBC clonal dynamics

Since SAG induced dynamic changes in the stem cell pool over a short period (Fig. 2), we next investigated the effects of SAG on Lgr5^+^ CBC clonal dynamics. We traced Lgr5^+^ CBC clones using *Lgr5-tdTomato* mice, in which tamoxifen treatment leads to the labeling of Lgr5^+^ CBC descendants with tdTomato (Fig. 3a-d). In tamoxifen-injected mice, tdTomato-labeled clones reached the middle of the intestinal villi by dpi 3 and the tips of the villi by dpi 7. In contrast, when we co-injected mice with tamoxifen and SAG, most of the tdTomato-labeled clones had already reached the tips of the villi by dpi 3 (Fig. 3c). Notably, although SAG accelerated clonal expansion up to dpi 3, these SAG-induced clones did not maintain dominance to dpi 7. Instead, we observed a restriction of tdTomato expression to regions near the crypts on dpi 7 (Fig. 3c).

**Fig. 3.**
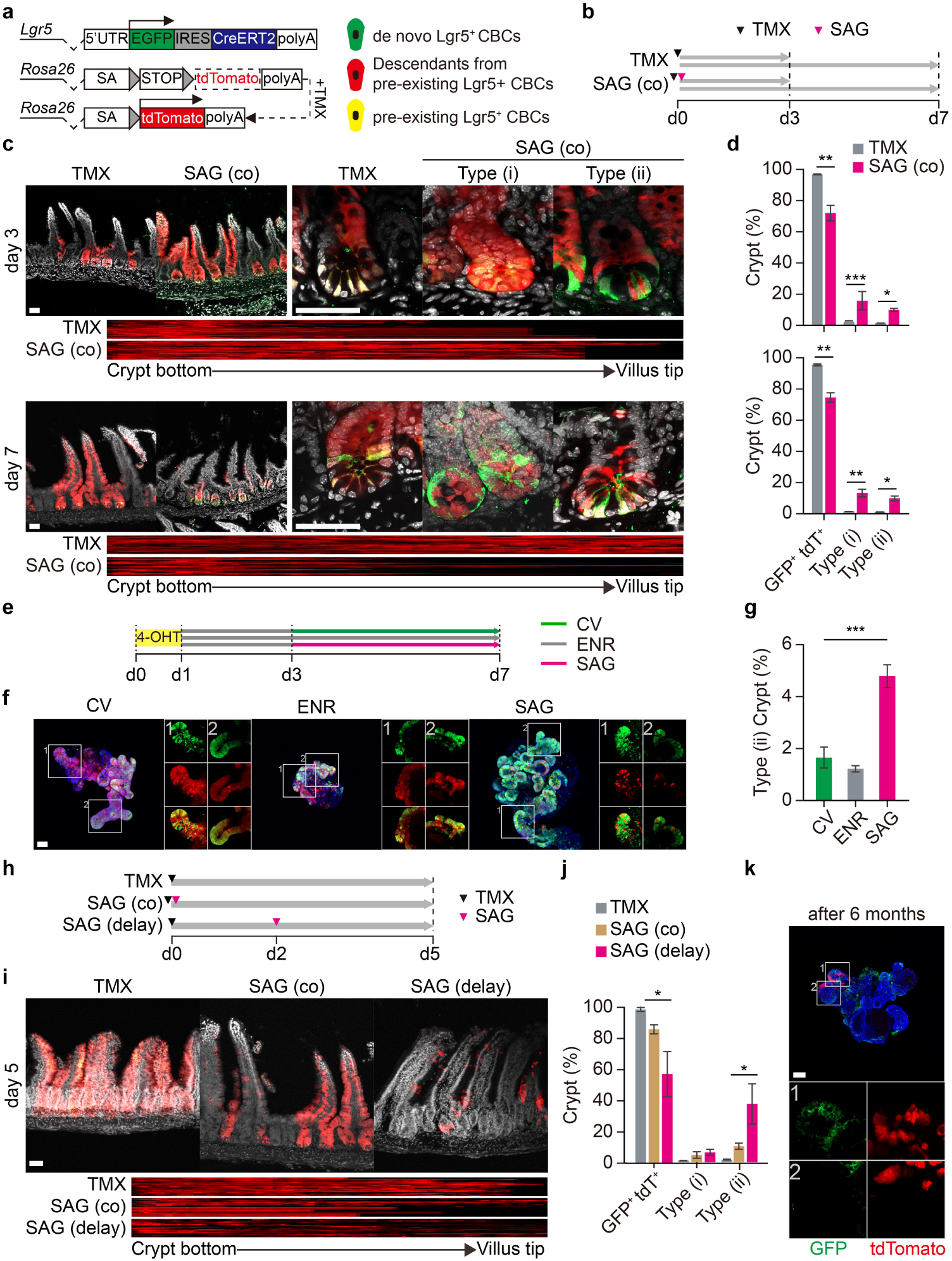
SAG triggers production of de novo Lgr5^+^ CBCs and facilitates stem cell dynamics. **a**, Genetic schematic for the lineage tracing of Lgr5^+^ CBCs (left). TMX injection produces three different patterns of fluorescent labeling: de novo Lgr5^+^ CBCs (green), cells descending from pre-existing Lgr5^+^ CBCs (red), and pre-existing Lgr5^+^ CBCs (yellow). **b**, Timeline for TMX and SAG co-administration (**c**, **d**). **c**, Representative confocal images of crypt-villus units on day 3 or day 7 after treatment with TMX or SAG. Type (i): crypts with tdTomato^+^ Paneth cells (PCs). Type (ii): crypts with de novo Lgr5^+^ CBCs. Heatmap showing the distribution of tdTomato labeling along the crypt-villus axis. *n* = 5 mice per group. **d**, Proportion of GFP^+^ tdTomato^+^, type (i) and type (ii) crypts on day 3 (*n* = 74, 82) and day 7 (*n* = 94, 138) **e**, Timeline for (**f**, **g**). 4-OHT, 4-hydroxytamoxifen. **f**, Representative confocal images showing GFP and tdTomato expression in CV, ENR, and SAG-treated organoids. In the right insets, crypt regions are magnified and channel-split. **g**, Proportion of type (ii) crypts in organoids. (*n* = 30, 31, 33). **h**, Timeline for the delayed injection of SAG (**i**, **j**). **i**, Representative confocal images of crypt-villus units on day 5 after treatment with TMX. Heatmap showing the distribution of tdTomato labeling along the crypt-villus axis. **j**, Proportion of GFP^+^ tdTomato^+^, type (i) and type (ii) crypts (*n* = 135, 80, 141). **k**, Representative confocal images showing long-retained tdTomato-expressing cells in organoids from SAG-delayed mice cultured for 6 months in vitro. Organoid experiments were performed at least in triplicate. Scale bar, 50 µm. Two-tailed Mann–Whitney U test (**d**, **g**, and **j**). \**P* < 0.05, \*\**P* < 0.01, and \*\*\**P* < 0.001.

In magnified crypt images from tamoxifen-injected mice, we observed Lgr5^+^ CBCs and their descendant cells labeled by tdTomato with unlabeled PCs positioned between the Lgr5^+^ CBCs at both dpi 3 and dpi 7. Surprisingly, we observed two novel tracing patterns in mice injected with both SAG and tamoxifen that we never observed in mice injected only with tamoxifen. In the first pattern (i), we found tdTomato labeling of PCs as well as Lgr5^+^ CBCs on dpi 3 (Fig. 3c, d). Considering the long lifespan of PCs (∼60 days)^36^, this suggests accelerated PC turnover. Even more strikingly, the second pattern (ii) included a significant proportion of Lgr5^+^ CBCs that lacked tdTomato labeling (Fig. 3c, d). Since most IECs are typically derived from Lgr5^+^ CBCs, these tdTomato-negative Lgr5^+^ CBCs strongly indicate that SAG treatment triggers the production of de novo Lgr5^+^ CBCs independently from pre-existing Lgr5^+^ CBCs.

Next, we reproduced our Lgr5^+^ CBC lineage tracing experiment in vitro using organoids derived from *Lgr5-tdTomato* mice (Fig. 3e-g). In ENR- and CV-treated organoids additionally treated with 4-hydroxytamoxifen (4-OHT), we observed tdTomato expression in most Lgr5^+^ CBCs by day 7. In contrast, in organoids treated with SAG from day 3, we observed numerous Lgr5^+^ CBCs lacking tdTomato expression (Fig. 3f, g). Thus, our organoid tracing data are consistent with our in vivo lineage tracing data. Together, these findings indicate Hh signaling activation both accelerates cellular turnover in existing clones and triggers the de novo generation of Lgr5^+^ CBCs in a mechanism unrelated to their self-replication in vivo.

To gain more insight into the effects of SAG, we next injected mice first with tamoxifen and then again with SAG two days later (Fig. 3h-k). When we compared these mice with mice treated simultaneously with tamoxifen-SAG, we found an exacerbation of the abnormal crypt patterns and an even stronger attrition of the tdTomato clones (Fig. 3i, j). Consequently, most of the remaining tdTomato-labeled cells were restricted to the upper crypt and isthmus regions (Fig. 3i, j, Extended Data Fig. 4c, d). Although these residual tdTomato-labeled cells were long-retained, including both Lgr5^+^ CBCs and Lgr5^-^ cells, they no longer produced descendant clones in the villi. Strikingly, when we cultured intestinal organoids from the crypt clumps of these mice and passaged them as dissociated clumps through several passages, the tdTomato-labeled cells remained near the organoid crypt regions even after 6 months of culture (Fig. 3k). These findings suggest Hh signaling activation drives some existing stem cells and their clones into a quiescent state.

### SAG and PGE_2_ induce distinct regenerative mechanisms in crypts and villi

Next, we wondered whether Hh activation is part of the recently proposed dedifferentiation mechanism or some other unexplored mechanism. We compared organoids cultured with ENR, SAG, or prostaglandin E_2_ (PGE_2_) (Fig. 4a) because PGE_2_ rapidly induces organoids to undergo a fetal-like transformation via dedifferentiation^11,12^. Although SAG induced crypt changes (Fig. 2), PGE_2_ rapidly expanded villi while also preventing cryptic budding (Fig. 4b and Extended Data Fig. 5a, b). Interestingly, co-administration of SAG and PGE_2_ resulted in separate populations of villi-expanded and crypt-enlarged organoids, with none simultaneously exhibiting both phenomena (Extended Data Fig. 5a, b). In IV-treated organoids, which show EC enrichment, PGE_2_ still caused expanded villi, whereas SAG did not cause crypt enlargement (Extended Data Fig. 5c, d). These data indicate that PGE_2_ acts primarily on villi, while SAG acts primarily on crypts.

**Fig. 4.**
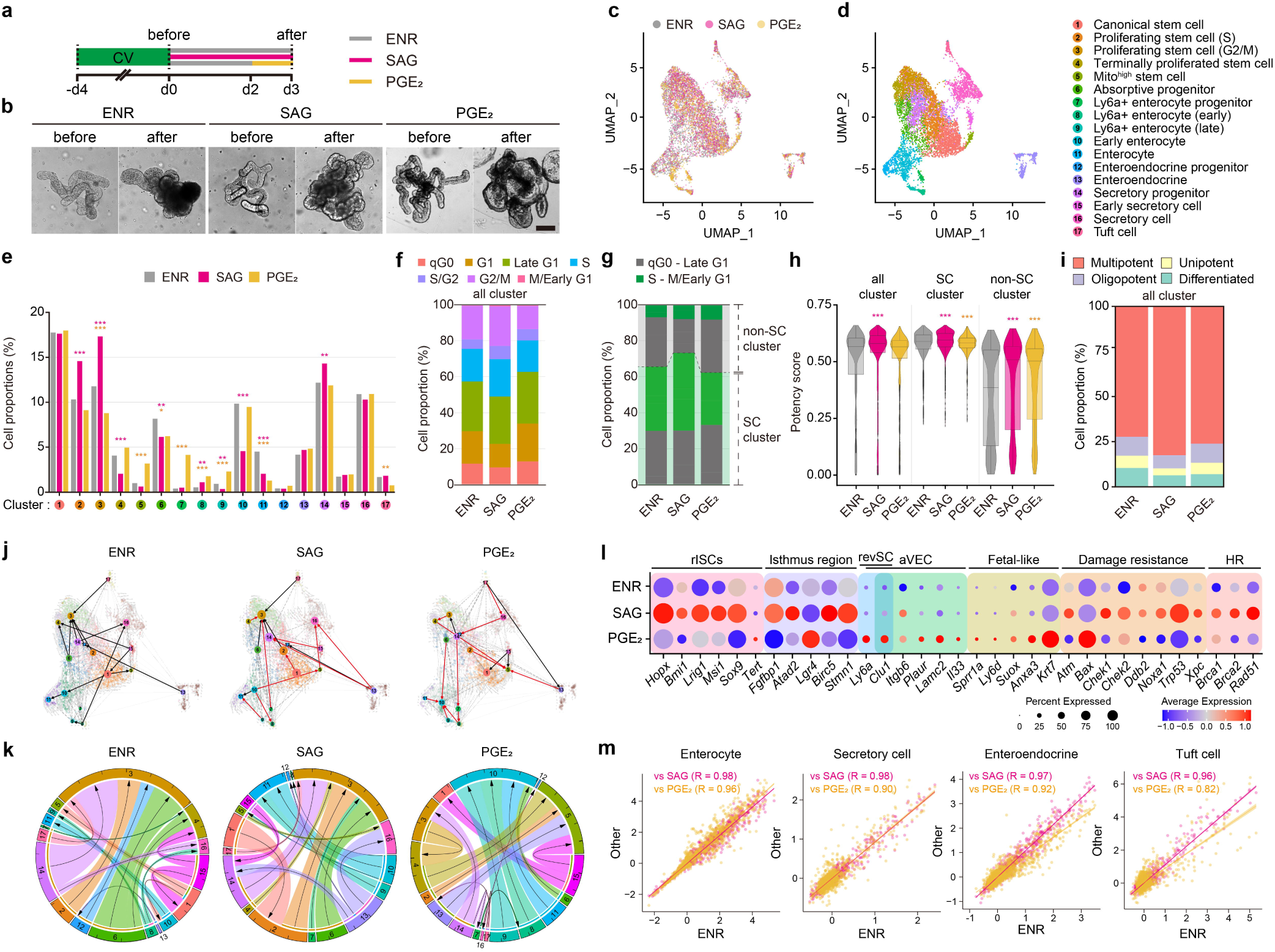
SAG promotes the proliferation of stem cells, whereas PGE2 induces fetal-like dedifferentiation. **a**, Experimental timeline. **b**, Representative brightfield images showing morphological changes in organoids before and after treatment. Scale bar, 50 μm. **c**, UMAP plot of single-cell transcriptomes colored according to culture condition (ENR, SAG, PGE□). **d**, UMAP plot of single-cell transcriptomes separated into colored clusters 1-17, as defined by lineage identity (stem, secretory, absorptive) and cell cycle state. **e**, Bar plots showing the proportion of cells from each culture condition group across the 17 clusters. **f**, Stacked bar plots showing the proportion of cells in distinct cell cycle phases (qG0, G1, Late G1, S, S/G2, G2/M, M/Early G1) across treatment conditions. **g**, Comparison of low-proliferative (qG0–Late G1) and actively cycling (S–M/Early G1) populations within stem cell (SC) versus non-stem cell (non-SC) clusters. **h**, Violin plots showing CytoTRACE2 potency scores across culture conditions for all clusters, SC clusters, and non-SC clusters. **i**, Stacked bar plots showing the proportion of multipotent, oligopotent, unipotent, and differentiated cells in each culture condition according to their CytoTRACE2 classifications. **j**, PAGA and scVelo analyses highlighting lineage connectivity and differentiation directionality under each treatment condition; red arrows denote differentiation trajectory deviations compared to the ENR baseline. **k**, Chord diagrams reconstructed from PAGA and scVelo-inferred transitions (**j**), summarizing directional lineage connectivity between clusters under each condition. Tick marks reflect the relative magnitude of transitions, displayed in increments of 0.2. **l**, Dot plot showing the expression of representative genes across culture conditions associated with reserve ISCs (rISCs), the isthmus region, revival stem cells (revSC), atrophy-induced villus epithelial cells (aVEC), fetal-like states, damage resistance, and homologous recombination (HR). Dot size reflects the proportion of cells, and dot color indicates average expression level. **m**, Pearson correlation plots of pseudobulk gene expression for 2,000 highly variable genes, comparing ENR and SAG or PGE□ treatments in mature differentiated cell types. \**P* < 0.05, \*\**P* < 0.01, and \*\*\**P* < 0.001.

To identify the mechanisms underlying the cell type-specific differences in the effects of SAG and PGE_2_ treatment, we conducted single-cell RNA sequencing (scRNA-seq) on organoids cultured under the ENR, SAG, or PGE_2_ conditions. We collected biological replicates for each condition and obtained a total of 8,976 cells (ENR: 3,819 cells; SAG: 3,285 cells; PGE_2_: 1,872 cells) that passed our quality control criteria. We then visualized the data using the uniform manifold approximation and projection (UMAP) and applied graph-based algorithms. Using canonical cell-type specific markers, we identified a total of 17 distinct clusters (Fig. 4c, d, and Extended Data Fig. 6a, b). Four of these were typical, mature differentiated cells, namely enterocytes, enteroendocrine cells (EECs), tuft cells, and secretory cells with mixed features of both goblet cells and PCs. The remaining clusters included two clusters each of early-state enterocytes and secretory cells, three Ly6a^+^ enterocyte-related lineages, and stem/progenitor cell populations. To further dissect stem cell dynamics, we divided the stem/progenitor compartment into eight subgroups, including canonical stem cells, proliferating cells in the S and G2/M phases, terminally proliferated cells, absorptive progenitors, secretory progenitors, EEC progenitors, and Mito^high^ stem cells, each harboring distinct cell cycling states estimated by ccAFv2 analysis (Extended Data Fig. 7a, b), and specific makers for stem cell subpopulations previously identified in public data (GSE148524)^37^.

Compared with control organoids, those treated with SAG and PGE_2_ showed distinct changes in both cell-type composition and cell state. SAG treatment significantly expanded the proportion of proliferating cells in the S and G2/M phases and increased secretory progenitors, while reducing terminally proliferated cells and the enterocyte-related lineages (Fig. 4e). This shift increased actively cycling cells (i.e., those in S to M phase) both in the total population and specifically within the stem/progenitor compartment, while also decreasing non-stem/progenitor cells in qG0 or G1 phase (Fig. 4f, g). Moreover, SAG treatment increased CytoTRACE2 scores in both stem/progenitor and non-stem/progenitor cells, indicating an enrichment of multipotent cells (Fig. 4h, i and Extended Data Fig. 7c, d). RNA velocity and PAGA analyses further revealed that SAG treatment reorganized differentiation trajectories and inter-cluster connectivity patterns (Fig. 4j, k). The most prominent change was an intensified directional flow toward G2/M-phase proliferating cells. In addition to triggering the emergence of new direct trajectories from terminally proliferated cells and Ly6a□ enterocyte progenitors to G2/M proliferating cells, SAG also enhanced the directional flow toward secretory progenitors and S-phase proliferating cells, which serve as the primary upstream contributors to G2/M-phase proliferating cells.

These findings suggest SAG treatment orchestrates more efficient coordination of cell cycle progression. Thus, our results demonstrate that Hh signaling promotes stem/progenitor pool expansion by accelerating cell cycle progression while also maintaining cell-type diversity.

In contrast, PGE_2_ treatment increased the proportion of Ly6a^+^ enterocyte-related lineages and Mito^high^ stem cells while reducing normal enterocytes (Fig. 4e). Unlike SAG, PGE_2_ exerted minimal effects on cell cycle dynamics, causing only a slight decrease in the proportion of cells in G2/M phase (Fig. 4f, g). It also caused a modest increase in CytoTRACE2 scores for the non-stem/progenitor compartments (Fig. 4h, i). Trajectory analyses revealed that PGE_2_ strengthened directional flow toward early enterocytes and Mito^high^ cells while simultaneously weakening the flow toward G2/M proliferating cells (Fig. 4j).

Next, pseudobulk RNA-seq analysis further highlighted contrasting transcriptional responses underlying IEC regeneration between the two treatments. SAG treatment upregulated genes characteristic of rISCs^3-10^, isthmus stem cells^38,39^, DNA damage resistance, and homologous recombination pathways. In contrast, PGE_2_-treated organoids showed marked enrichment for genes associated with revSCs^40,41^, aVECs^41,42^, and fetal-like IECs^43^ (Fig 4l, Extended Data Fig. 8a-d). Despite showing enhanced proliferation, SAG-treated organoids did not exhibit significant changes in gene expression within the typical mature differentiated cell clusters compared to ENR-treated organoids (Fig. 4m). This suggests that Hh activation preserves the identify and functional quality of differentiated cell types while inducing transcriptional program shifts that reflect a higher proportion of cells actively engaged in DNA replication and repair, particularly homologous recombination. In contrast, PGE_2_-treated organoids induced significant transcriptional changes within mature differentiated cell populations.

Collectively, our scRNA-seq and organoid analyses delineated two fundamentally distinct regenerative mechanisms. While SAG facilitates regeneration by accelerating cell cycle progression within the stem/progenitor compartment, PGE_2_ drives dedifferentiation of enterocyte-related lineages.

### Hh signaling promotes damage resistance among IECs

Since SAG enhanced rISC- and isthmus stem cell-like features and damage resistance-related genes in our scRNA-seq data (Fig. 4l), we next asked whether Hh activation enhances IEC damage resistance. To test this, we compared the survival of organoids treated with ENR, SAG, or CV following radiation-induced damage (6 Gy) (Fig. 5a). At rec 5, SAG-treated organoids and CV-treated organoids exhibited the highest and lowest survival rates, respectively (Fig. 5b, c). Among the surviving organoids, those treated with SAG were larger in diameter than the others. Daily tracking of individual organoids revealed a progressive loss of pre-existing crypts, accompanied by the emergence of new crypts. While most pre-existing crypts in ENR- and CV-treated organoids were lost, SAG-treated organoids retained a subset of these crypts (Fig. 5e). Furthermore, SAG treatment significantly enhanced de novo crypt formation compared to other treatments (Fig. 5e, Extended Data Fig. 10). Notably, we found increased damage resistance when SAG was administered to ENR-, CV-, or CD-treated organoids, but not to IV-, ID-, or DV-treated organoids (Extended Data Fig. 9a-f). This suggests the involvement of stem cells within crypt regions, rather than those in the villi or PCs.

**Fig. 5.**
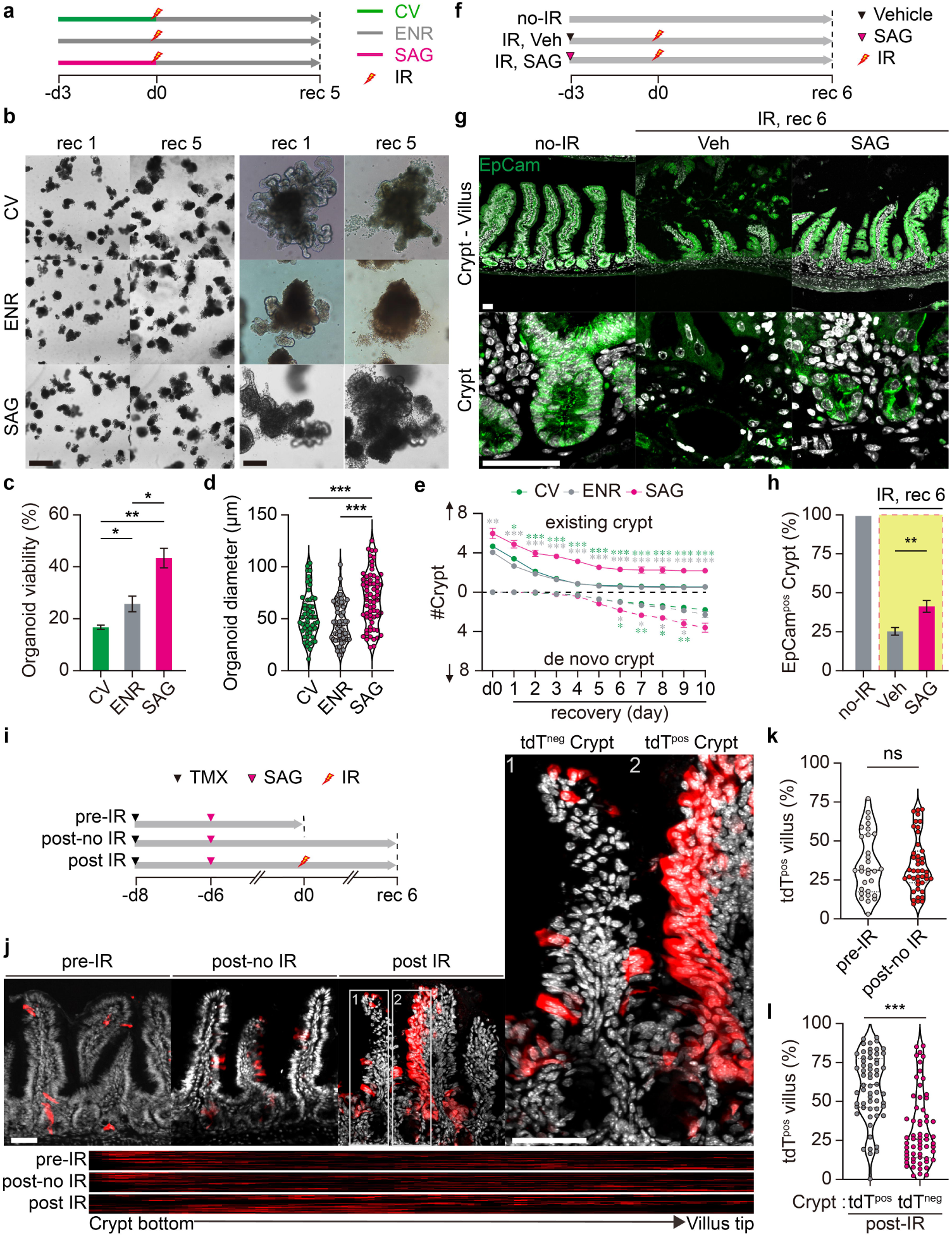
SAG-induced long-retained cells generate damage-resistant clones. **a**, Timeline for the organoid irradiation experiment (**b**-**d**). IR, irradiation. **b**, Representative bright field images of organoids cultured under the indicated conditions on recovery days 1 and 5. **c**, Organoid viability on dpi 5. **d**, Organoid diameter on dpi 5. (*n* = 56, 65, 65). **e**, Number of existing and de novo crypts arising by dpi 10 (*n* = 56, 51, 32). All these organoid experiments were replicated five times. **f**, Timeline for the in vivo irradiation experiment (**g**, **h**). veh, vehicle. **g**, Immunofluorescent images of small intestine stained with anti-EpCam (green). **h**, Proportion of EpCam^+^ crypts. (*n* = 51, 59, 65, from each of 3 mice). **i**, Timeline for irradiating long-retained tdTomato^+^ cells in vivo (**j**-**l**). **j**, Representative confocal images of the small intestine from pre-IR, post-noIR, and post-IR mice. In the post-IR images, insets are magnified on the right. Heatmap showing the distribution of tdTomato labeling along the crypt-villus axis. **k**, Relative occupancy of tdTomato^+^ cells in the villi of pre-IR and post-noIR mice. (*n* = 30, 43 from each of 4 mice). **l**, Relative occupancy of tdTomato^+^ cells in the villi of tdT^pos^ and tdT^neg^ crypts in post-IR mice. (*n* = 60, 61 from each of 4 mice). Scale bars, 50 µm. Unpaired two-tailed *t*-test (**c**), ANOVA or Kruskal–Wallis test (**d**), mixed-effects model followed by Tukey’s multiple comparisons test (**e**), one-way ANOVA with Tukey’s test (**h**), and Mann–Whitney test (**k, l).** \**P* < 0.05, \*\**P* < 0.01, and \*\*\**P* < 0.001.

Next, we exposed vehicle- and SAG-injected mice to a sublethal dose of radiation (6 Gy) and examined histologic changes to the intestinal epithelium (Fig. 5f-h). In vehicle-injected mice, irradiation led to detachment of the villus epithelium from the underlying mesenchyme, reduction in EpCam^+^ crypts, and increased expression of the apoptotic marker CC-3. In contrast, SAG-injected mice exhibited relatively intact epithelial layers with reduced villus detachment, lower CC-3 expression, and better preservation of EpCam^+^ crypts. (Fig 5g, h and Extended Data Fig. 9g-i). These data suggest Hh signaling confers damage resistance upon IECs.

Finally, we sought to determine whether the quiescent and long-retained tdTomato-labeled crypt cells induced by SAG treatment contribute to damage resistance. To produce these residual tdTomato-labeled cells in *Lgr5-tdTomato* mice, we injected the mice with SAG two days after tamoxifen injection (Fig. 5i). Then, six days after SAG injection (in pre-IR mice), we found tdTomato-labeled cells restricted to the crypts and isthmus regions along with de novo Lgr5^+^ CBCs in the crypts, showing minimal expansion into the villi. Notably, we observed very few crypt-villus units continuously occupied by tdTomato-labeled clones, and only a few scattered tdTomato-labeled cells in the villi (Fig. 5j-i). Six days after irradiation (post-IR mice), however, we observed two distinct tdTomato-labeling patterns in the crypt-villus units. Where tdTomato-labeled cells remained in the crypts and isthmus, the connected villi were continuously occupied by tdTomato-labeled clones. In contrast, when tdTomato-labeled cells were absent, we also observed reduced crypt-villus unit occupancy (Fig. 5j-i). These findings indicate that residual tdTomato-labeled cells within the crypts undergo clonal expansion upon damage, ruling out the possibility of villus cell dedifferentiation. In non-irradiated mice harvested at the same time as post-IR mice (post-noIR mice), tdTomato labeling remained confined to the crypt and isthmus regions without expansion into the villi, like pre-IR mice (Fig. 5j-i). These results demonstrate that the changes we observed in post-IR mice were specifically induced by radiation-induced damage rather than merely resulting from time progression.

Together, these data suggest Hh signaling activation induces dormant cells in the crypt and isthmus regions that do not normally generate dominant clones in crypt-villi units under normal conditions but subsequently expand to generate clones in response to radiation damage. This is reminiscent of the concept of intestinal reserve stem cells.

## Discussion

Based on our findings, we propose a model in which Hh signaling plays a central role in the regeneration of IECs. In the short term, Hh signaling promotes ISC clone competition characterized by accelerated cellular turnover and the generation of de novo Lgr5^+^ CBCs (Fig. 3). This process leads to the extinction/attrition of previously dominant lineages, while also allowing stem cells that had previously remained in reserve to acquire clonal dominance. Meanwhile, other competition-recessive stem cells remain dormant within the crypts and isthmus (Fig. 3). In the long term, these dormant cells exhibit resistance to subsequent damage (Fig. 5), whereas the dominant clonal lineages are susceptible, enabling the continuous replacement of lineages in response to damage. Our model not only supports the previous hypothetical model that posited reserve stem cells, but it also provides a possible explanation for how the reserve stem cells are awakened and generated. According to recent reports focused on isthmus stem cells^38,39^, the de novo Lgr5^+^ CBCs we observed may be derived from Fgfbp1^+^ or Lgr4^+^ isthmus stem cells.

In our comparative investigation, we have clearly delineated complementary roles for Hh and PGE_2_ signaling in IEC regeneration. Hh signaling targets crypt and isthmus stem cells, promoting the selection of healthy, damage-resistant clones. This is a fundamental strategy that permits a complete reconstruction of crypt-villi units, but it takes considerable time to reach distal end villi. To bridge the transient gap between injury and Hh signaling-induced reconstruction, PGE_2_ signaling can rapidly facilitate villi expansion by acting primarily on ECs and their progenitor cells, providing temporal repair. Due to their mutually exclusive target regions and temporal kinetics (Fig. 4), these multilayered repair mechanisms may be the fundamental basis for the exceptionally superior tissue homeostasis of the intestinal epithelium.

In the intestine, Hh signaling has been extensively studied for its developmental roles in generation of enteric neurons^25^ and smooth muscle layers^25^, as well as in villus formation and patterning^27,28^. In adults, despite its diminished contribution, Hh signaling still plays a role in diseases such as gut inflammation^32,44^ and cancer^29,45^. Most studies have posited a paracrine signaling mechanism in which Hh ligands secreted by epithelial cells act on the mesenchymal compartment, regulating mesenchymal cell behavior and inducing feedback loops that influence epithelial cells^26,27,29,32,44,45^. Our findings, however, challenge this paradigm by demonstrating that Hh signaling can directly regulate ISC behavior in adults. Using organoids composed solely of IECs (Fig. 2), we found that pharmacologic manipulation of Hh signaling altered ISC activity, implying IEC expression of Hh receptors. In a reanalysis of public scRNA-seq data from intestinal organoids (GSE148524), not only did we find abundant expression of Hh ligand (*Ihh*) in ECs and PCs^46^, but we also found expression of the Hh signaling receptors *Smo* and *Ptch1* in ISC subsets (Extended Data Fig. 7h). Additionally, recent studies have demonstrated the presence of primary cilia—the subcellular organelles responsible for Hh signaling transduction—in BMI1^+^ intestinal crypt epithelial cells^47^. Given that primary cilia form during the quiescent G0 phase, it is plausible that quiescent ISCs overlap with epithelial cells capable of responding to Hh signaling. It is thus possible that previous studies have primarily focused on mesenchymal responses because such a limited proportion of epithelial cells respond to Hh signaling. We acknowledge that our SAG-induced results demonstrate the maximal effects of Hh signaling under experimental conditions because SAG can activate Smoothened independently of primary cilia^48^. It is likely that under physiological conditions, Hh signaling exerts more delicate effects, particularly on quiescent ISCs with primary cilia. Furthermore, direct epithelial Hh responses and indirect mesenchymal feedback may act synergistically to regulate ISC behavior in vivo. Further studies will be necessary to elucidate the role of primary cilia in the context of IEC regeneration and to determine how epithelial and mesenchymal components integrate Hh signaling to maintain ISC function and intestinal homeostasis.

Most previous studies of ISC regeneration relied on genetic lineage tracing using tamoxifen-inducible Cre drivers^3-7,16,18-20,40^. These drivers have also shown their ability to give rise to clonal labeling after Lgr5^+^ CBC injury. While powerful, this technique is inherently dependent on the target gene marker it employs. If the marker exhibits transient or leaky expression, even at extremely low levels, it can easily distort the identities of the cellular origins of traced clones. Consistent with recent reports^38,39^, we did not find any cell clusters annotated as reserve stem cells, nor could we find their signature genes in scRNA-seq data. Instead, we have clearly demonstrated that Hh signaling significantly alters cell cycle dynamics and differentiation trajectories. Not only do our findings help explain why this debate regarding the identity of the cells and the underlying mechanisms responsible for IEC regeneration has persisted for so long, they also offer a path toward its resolution. Rather than identifying specific gene markers for regenerative cell populations, our findings highlight a signaling pathway that switches ISCs between dominant and quiescent states. Future investigations should focus on clarifying the interactions between Hh signaling and other regenerative pathways in human intestinal tissues and exploring potential therapeutic applications.

## Methods

### Animals

All animal procedures were conducted according to guidelines and protocols approved by the Institutional Animal Care and Use Committee of Korea University College of Medicine (approval number: KOREA-2024-0015). *Lgr5-eGFP-IRES-CreERT2* (#JAX008875), *Rosa26-lsl-tdTomato* (#JAX007908), and *Rosa26-lsl-DTA* (#JAX009669) mice were obtained from the Jackson Laboratory. *Lgr5-tdTomato* and *Lgr5-DTA* mice were generated by breeding *Lgr5-eGFP-IRES-CreERT2* to *Rosa26-lsl-tdTomato* or *Rosa26-lsl-DTA* mice, respectively. All mice were maintained under standard conditions with a 12-hour light/dark cycle, controlled temperature (19–23°C) and humidity, and ad libitum access to food and water.

### Organoids culture

#### Conditioned media (CM)

RSPO1 CM was generated from an R-Spondin1-expressing 293T Cell Line (SCC111, Merck, Darmstadt, Germany). Noggin CM was generated from a Noggin-expressing stable cell line, generously gifted by Dr Peihua Jiang (Monell Chemical Senses Center). The cell lines were maintained and selected in Dulbecco’s Modified Eagle Medium (DMEM; 11995-065, Thermo Fisher Scientific, Waltham, MA, USA) supplemented with 10% fetal bovine serum (FBS; 12483020, Thermo Fisher Scientific) and 400 µg/mL Zeocin (R-25001, Thermo Fisher Scientific). After at least two days of selection, the cells were washed with DMEM without Zeocin and subsequently cultured in Advanced DMEM/F12 (12634010, Thermo Fisher Scientific) with 1% GlutaMax (35050-061, Gibco, Waltham, MA, USA) for 10 days to produce RSPO1 CM or Opti-MEM (31985070, Thermo Fisher Scientific) for 7 days to produce Noggin CM, respectively. Both CM were collected, centrifuged at 2000 x g for 15 minutes at 4°C, sterile filtered, and stored at −80°C for up to 2 months.

#### Crypt isolation and organoid culture

Crypt epithelial cells were released by mechanical agitation and filtered through a 70 μm cell strainer (93070, SPL, Pocheon, Korea). The isolated crypts were mixed with Matrigel (354230, Corning, NY, USA) at a 1:1 ratio in basal media. A total volume of 50 μL per well was added to the center of each well of a 24-well plate (30024, SPL). Basal media consisted of Advanced DMEM/F12 supplemented with 2 mM GlutaMax, 100 U/mL Penicillin-Streptomycin, 1% N-2 supplement (17502-048, Gibco), and 2% B-27 supplement (17504-044, Gibco). This basal media was used for organoid washing, passaging, and as a base for the organoid culture media. To prepare organoid culture media or ENR, 80% basal media was supplemented with 10% Noggin CM, 10% RSPO1 CM, 50 ng/mL EGF (AF-100-15-A, PeproTech, Rocky Hill, NJ, USA), and 1 mM N-acetylcysteine (A9165, Sigma-Aldrich, St Louis, MO, USA).

#### Passaging

For each passage, organoids cultured for 7 days were harvested by filling the Matrigel dome with cold phosphate-buffered saline (PBS), followed by washes with basal media. The organoids were then dissociated into cell clumps by pipetting several times, mixed with basal media, and split 1:4. Passaged organoid cell clumps were embedded in a 1:1 mixture of Matrigel and media, plated in the center of each well of a 24-well plate (200–300 organoids per well), and expanded in a 3-day CV start-culture. CV start-culture media was prepared by adding 3 μM CHIR-99021 (SML1046, Sigma-Aldrich) and 1 mM valproic acid (P4543, Sigma-Aldrich) to ENR culture media. CV start-culture media supports the self-renewal of *Lgr5*^+^ CBCs, leading to the production of a homogenous pool of *Lgr5*^+^ CBCs at the beginning of the experiments.

### In vivo experimental designs (chemical administration and irradiation)

#### Chemical stocks

Tamoxifen (GC17901, GLPbio, Montclair, CA, USA) was dissolved in corn oil (C8267, Sigma-Aldrich). 4-hydroxytamoxifen (4-OHT; H7904, Merck) was dissolved in dimethyl sulfoxide (DMSO; D2650, Sigma-Aldrich). SAG (HY-12848, MedChemExpress, Monmouth Junction, NJ, USA) was dissolved at 0.5 mg/ml in DMSO to make a stock and then further dissolved at 2 mg/ml in PBS for use. Cyclopamine was first prepared by mixing hydroxypropyl-β-cyclodextrin (HBC; H107, Sigma-Aldrich) in 1 ml PBS. Then, the cyclopamine was suspended in 45% HBC solution and allowed to mix for 24 hours at room temperature to ensure complete dissolution.

### In vivo drug administration

For the Lgr5^+^ CBC ablation experiments (Fig. 1 and Extended Data Fig. 1), *Lgr5-DTA* mice and *Lgr5-eGFP-IRES-CreERT2* genetic controls were subjected to intraperitoneal injections of tamoxifen at 10 mg/kg/day for five consecutive days. For the tamoxifen-cyclopamine co-administration groups, *Lgr5-DTA* mice received separate bolus injections of cyclopamine (5 mg/kg). For the bulk RNA-seq experiments, mice were harvested the day after their final injection. For the immunohistochemistry experiments, mice were harvested seven days after their final injection. For the SAG injection experiments (Fig. 2), *Lgr5-eGFP-IRES-CreERT2* mice received a single intraperitoneal injection of SAG at 10 mg/kg before being sacrificed at the indicated time points. For the lineage tracing experiments (Fig. 3 and Extended Data Fig. 4), *Lgr5-tdTomato* mice received a single intraperitoneal injection of tamoxifen before being sacrificed at the indicated time points. In the SAG co-administered groups (Fig. 3b-d, Extended Data Fig. 4a, b), SAG was additionally administered via separate bolus injection on the same day as the tamoxifen injections. In the delayed SAG administration group (Fig. 3h-k, Extended Data Fig. 4c, d), SAG was injected two days after the tamoxifen injections. For the in vivo irradiation experiments (Fig. 5f-h and Extended Data Fig. 9g-i), B6N mice were injected with either vehicle or SAG three days before radiation exposure. The mice were then sacrificed six days after irradiation. For the combined lineage tracing and irradiation experiments (Fig. 5i-l and Extended Data Fig. 9j), *Lgr5-tdTomato* mice received a single intraperitoneal injection of tamoxifen eight days before radiation exposure and another of SAG six days before radiation exposure. Pre-IR mice were sacrificed on the day of irradiation. Post-IR mice were sacrificed six days after radiation exposure. Post-noIR mice were sacrificed on the same day as post-IR mice but without the radiation exposure.

### In vitro experimental designs (chemical administration and irradiation)

For the in vitro experiments (Fig. 2e-i, Extended Data Fig. 3b), crypt epithelial cells were isolated from *Lgr5-eGFP-IRES-CreERT2* mice and cultured into organoids. After several passages, the organoids were initially cultured under CV conditions for 4 days, and then divided into three groups (i.e., CV, ENR, and SAG). For the SAG group, 1 μM SAG was added to ENR culture media, and then the organoids were cultured for an additional 3 days. For the holotomographic time-lapse imaging experiment (Extended Data Fig. 3b), organoids pretreated under CV conditions were divided into three groups (i.e., CV, ENR, and SAG). Images were captured immediately after the organoids were treated with CV, ENR, or SAG. For the organoid lineage tracing experiments (Fig. 3e-g), crypt epithelia were isolated from *Lgr5-tdTomato* mice and cultured into organoids. Organoids were treated with 4-OHT for one day and then washed and re-seeded into fresh Matrigel to eliminate residual 4-OHT. After three days of culture in ENR medium, the organoids were treated with CV, ENR, or SAG. To minimize potential mesenchymal cell contributions, lineage tracing was performed using the organoids subjected to at least two passages in vitro. For the scRNA-seq experiment (Fig. 4, Extended Data Figs. 6, 7, and 8), organoids precultured for four days under CV start conditions were divided into three groups. The ENR and SAG groups were cultured under the ENR or SAG condition for three days, respectively. The PGE_2_ group was maintained under the ENR condition for two additional days before being treated with PGE_2_ (1 μM, 2296, Tocris Bioscience, Bristol, UK) for a single day. For the in vitro irradiation experiments (Fig. 5a-e, Extended Data Figs. 9d, f and 10), organoids were cultured under CV, ENR, or SAG conditions for three days prior to irradiation. Following irradiation, the culture media was replaced with ENR, and morphological changes were monitored daily until recovery day 10. During this time, the media was refreshed every three days. For the in vitro experiments on organoids precultured under various conditions with SAG (Extended Data Fig. 9), organoids pre-cultured for three days under CV start conditions were divided into six groups (i.e., ENR, CV, CD, ID, IV, and DV). The organoids were maintained under each condition for two days. Following this, the organoids were either cultured under ENR or with SAG for an additional three days. In addition to being subjected to qRT-PCR, morphology was also traced daily until recovery day 6. For each condition, ENR media was supplemented with the following: CV with C (CHIR-99021; 3 μM, SML1046, Sigma-Aldrich) and V (Valproic acid; 1 mM, P4543, Sigma-Aldrich), CD with C and D (DAPT; 10 μM, HY-13027, MedChem Express), ID with I (IWP-2; 2 μM, I0536, Sigma-Aldrich) and D, IV with I and V, and DV with D and V.

### Irradiation

For in vivo irradiation (IR) exposure, anesthetized mice were subjected to 6 Gy whole-body IR using an RS-320 (X-Strahl Ltd, Walsall Wood, UK) in a fixed frame. For organoid IR exposure, culture plates containing organoid samples were irradiated with planar rotation to ensure uniform irradiation. After IR exposure, bright field images were captured daily to trace the morphological changes of each organoid.

### Immunohistochemistry

Mouse small intestines were harvested under anesthesia. The intestines were then cut longitudinally for Swiss rolling, fixed in 4% paraformaldehyde (PFA) in PBS for 6 hours, and then transferred to 30% sucrose in PBS overnight. The settled tissues were then embedded in Tissue-Tek OCT (4583, Sakura Finetechnical, Tokyo, Japan), frozen on dry ice, and stored at −80°C until further use. Cryoblocks were cut into 30 μm-thick sections on a cryostat (CM3050S, Leica Biosystems, Nussloch, Germany). The sections were then attached directly to slide glass (HMA-S9911, MATSUNAMI, Japan). The samples were permeabilized and blocked in 3% Triton X-100 in PBS (PBST) with 3% Bovine Serum Albumin (BSA; BSAS 0.1, Bovogen Biologicals, Keilor East, VIC, Australia) for 30 minutes at room temperature. The samples were then incubated with primary antibodies diluted in blocking solution overnight at 4°C. After three 10-minute washes in PBST, the samples were incubated with secondary antibodies for 1 hour at room temperature in wash buffer. After another three washes, VectaShield (H-1000, Vector Laboratories, Burlingame, CA, US) with DAPI (1:1000; 14285, Cayman Chemical, Ann Arbor, MI, USA) was added, and the samples were covered with glass coverslips.

For immunostaining with anti-EpCAM, the intestines were embedded in paraffin and cut into 5 μm-thick sections using a rotary microtome (RM2255, Leica Biosystems). Free-floating paraffin sections were placed onto charged glass slides. The slides were then deparaffinized in xylene and rehydrated through a graded ethanol series (100%, 95%, 70%) ending with deionized water. Antigen retrieval was conducted in citrate buffer (10 mM sodium citrate, 0.05% Tween 20, pH 6.0) at 121°C for 10 minutes. After the slides cooled to room temperature, the remaining antibody incubation procedures were performed as described above.

Intestinal organoids were harvested from Matrigel via gentle pipetting of cold PBS to dissolve the matrix. Organoid samples were then fixed in 4% PFA in PBS for 15 minutes and permeabilized at 0.2% PBST with 3% BSA for 1 hour at room temperature. Primary and secondary antibody incubations were performed as described above. Organoid samples were stained with DAPI and mounted on glass slides under Vectashield and coverslips. Confocal images were acquired using an LSM 900 (Zeiss, Oberkochen, Germany) or K1-Fluo (Nanoscopesystem, Daejeon, Korea) confocal microscope.

### Antibodies

The following primary antibodies were used: chicken polyclonal anti-GFP (GFP-1020, Aves Labs, Togard, OR, USA, RRID:AB_10000240), rabbit monoclonal anti-Tert (ab32020, Abcam, Cambridge, UK, RRID: AB_778296), rabbit monoclonal anti-Ki67 (ab16667, Abcam, RRID:AB_30245), rabbit monoclonal anti-cleaved Caspase-3 (9664, Cell Signaling Technology, Danvers, MA, USA, RRID:AB_2070042), rat monoclonal anti-Krt8 (TROMA-I, Development Studies Hybridoma Bank, Iowa City, IA, USA, RRID:AB_531826), rabbit monoclonal anti-EpCAM (ab223582, Abcam, RRID:AB_2762366), rabbit polyclonal anti-Lyz1 (PA1-29680, Thermo Fisher Scientific, RRID:AB_1955851), mouse monoclonal anti-Chga (sc-393941, Cruz Biotechnology, Dallas, TX, USA, RRID:AB_2801371), rabbit polyclonal anti-Muc2 (LS B16111, LSBio, Seattle, WA, USA), and rabbit polyclonal anti-Alpi (PA5-22210, Thermo Fisher Scientific, AB_11153471).

The following secondary antibodies were used: anti-chicken Alexa 488 (A32931, Invitrogen, Waltham, MA, USA, RRID:AB_2762843), anti-rabbit Alexa 488 (A32790, Invitrogen, RRID:AB_2762833), anti-rabbit Alexa 555 (A31572, Invitrogen, RRID:AB_162543), anti-mouse Alexa 555 (A32727, Invitrogen, RRID:AB_2633276), and anti-rat Alexa 555 (A48263, Invitrogen, RRID:AB_2896332).

### H&E staining

Paraffin-embedded intestines were cut into 5-μm thick sections. The sections were mounted on charged glass slides, deparaffinized in xylene (3 changes, 5 minutes each), and rehydrated via a graded ethanol series (100%, 95%, 70%). The sections were stained with hematoxylin (TA-125-MH, Thermo Fisher Scientific) for nuclear visualization, followed by eosin (786-1072, G-Biosciences, St Louis, MO, USA) to stain the cytoplasm. The stained sections were then dehydrated, cleared, and covered with Vectashield and glass coverslips.

### Quantitative real-time PCR

Total RNA was extracted from isolated crypt epithelial tissues or organoids using TRIzol (15596018, Thermo Fisher Scientific). Then, cDNAs were synthesized using a RevertAid RT Reverse Transcription Kit (K1622, Thermo Fisher Scientific). Quantitative qRT-PCR was conducted using SYBR Green qPCR Master Mix (GK10002, GLPbio) and the Quantstudio 3 (Thermo Fisher Scientific) according to the manufacturer’s instructions. Data analyzed using the delta-delta Ct (2^-ΔΔCt^) method were expressed as relative fold changes with normalization to *Gapdh or Actb* expression. The primers used were: *Lgr5-F*: 5’-CCTACTCGAAGACTTACCCAGT-3’; *Lgr5-R*: 5’-GCATTGGGGTGAATGATAGC-3’; *Olfm4-F*: 5’-AACATCACCCCAGGCTACAG-3’; *Olfm4-R:* 5’-TGTCCACAGACCCAGTGAAA-3’; *Tert-F*: 5’-AGCCGCACATTGGCTCTGCT-3’; *Tert-R*: 5’-TCGCCTCTGGGAGCTTCCGG-3’; *Gli1-F*: 5’-ACCACCCTACCTCTGTCTATTC-3’; *Gli1-R*: 5’-TTCAGACCATTGCCCATCAC-3’; *Gli2-F*: 5’-TGAAGGATTCCTGCCGTG-3’; *Gli2-R*: 5’-GAAGTTTTCCAGGACAGAACCA-3’; *Alpi*-F: 5’-GGTCAAGGCCAACTACAAGA-3’; *Alpi*-R: 5’-CACGGTACATCACTGAGAAGAC-3’; *Chga*-F: 5’-AGAACCAGAGCCCTGATGCCAA-3’; *Chga*-R: 5’-CTCTGTGGTTGCCTCAAAGCCA-3’; *Muc2*-F: 5’-CCATTGAGTTTGGGAACATGC-3’; *Muc2*-R: 5’-TTCGGCTCGGTGTTCAGAG-3’; *Lyz1*-F: 5’-GAGACCGAAGCACCGACTATG-3’; *Lyz1*-R: 5’-CGGTTTTGACATTGTGTTCGC-3’; *Actb-F*: 5’-CCACCAGTTCGCCATGGATGA-3’; and *Actb-R*: 5’-CATCACACCCTGGTGCCTAG-3’.

### Bulk RNA-sequencing

#### Sample preparation

Total RNA was extracted from isolated crypt epithelial tissues using TRIzol. RNA integrity was assessed using an Agilent 2100 Bioanalyzer (Agilent technologies, Santa Clara, CA, USA). Samples that achieved an RNA Integrity Number (RIN) > 7 were considered suitable for further processing. cDNA libraries were constructed using a SureSelect^XT^ RNA Direct kit (Agilent technologies). The resulting libraries were loaded onto an Illumina NovaSeq 6000 (Illumina, San Diego, CA, USA), and sequence data were generated using the recommended protocols.

#### Data analysis

Raw bulk RNA sequencing reads were aligned to the mouse reference genome (GRCm38) using HISAT2 version 2.2.1^29^. Quality control metrics, including junction saturation and read distribution across genomic features, were assessed using RSeQC version 5.0.1^5^. Transcript-level abundance quantification was performed against the GENCODE M34 reference transcriptome using Salmon version 1.10.0^49^. The resulting transcript abundance estimates were imported in R version 4.4.3 using tximport version 1.34.0^50^. Differential expression analysis was conducted using DESeq2 version 1.46.0^51^. P-values were adjusted for multiple comparisons using the Benjamini-Hochberg method^52^. Volcano plots were generated using EnhancedVolcano version 1.24.0^53^. Heatmaps were created using pheatmap version 1.0.12^54^. Gene Set Enrichment Analysis (GSEA)^55^ was conducted using fgsea version 1.32.4^56^ against the MSigDB^57^ and Gene Ontology (GO)^58^ databases. Functional enrichment was considered statistically significant at an adjusted p-value threshold of less than 0.05.

### Single-cell RNA sequencing

#### Sample preparation

Single-cell suspensions were prepared from ENR-, SAG-, and PGE_2_-treated organoid samples of more than 1 × 10□ cells per sample. The organoid samples were harvested and enzymatically dissociated into single cells. Briefly, the organoids were collected by centrifugation at 300 × g for 5 minutes and washed twice with cold PBS. The pellets were resuspended in TrypLE Express (12604013, Gibco) supplemented with actinomycin D (Act-D; final concentration 15 µM, A1410, Sigma-Aldrich) and DNase I (final concentration 100 U/mL, D5025, Sigma-Aldrich) to minimize transcriptional changes and reduce genomic DNA contamination. The suspensions were then incubated at 37°C for 2–3 minutes with gentle pipetting to promote cell dissociation. These reactions were stopped by adding basal medium and then subjected to centrifugation at 300 × g for 5 minutes. The cell pellets were then resuspended in PBS containing Act-D (15 µM) and DNase I (100 U/mL). Single-cell suspensions with > 80% viability, as determined by trypan blue exclusion, were used for downstream scRNA-seq experiments.

#### Single-cell library preparation and sequencing

Single-cell library preparation and raw data analysis were performed by SYSMED (Seoul, Korea). FACS-isolated single cells were counted using a LUNA-FL™ Dual Fluorescence Cell Counter (Logos Biosystems, Anyang, Korea). Cell samples from each culture condition were barcoded using the BD® Single-Cell Multiplexing Kit (BD Biosciences, Franklin Lakes, NJ, USA), which allowed us to combine 3 samples from different conditions and tissue types into a single pooled sample. The samples were then washed twenty times with staining buffer (2% FBS in PBS) using the Laminar Wash Mini1000 (Curiox Biosystems, Seoul, Korea). Pools of 20,000 cells were loaded on a microwell cartridge used with the BD Rhapsody Express system (BD Biosciences). Single-cell whole transcriptome libraries were prepared according to the manufacturer’s instructions using the BD Rhapsody WTA Amplification kit (BD Biosciences). The final libraries were quantified using a Qubit Fluorometer with the Qubit dsDNA HS Kit (Thermo Fisher Scientific). Their size distribution was measured using the Agilent high sensitivity DNA chip assay on a Tapestation system (Agilent technologies). Then, the generated single-cell libraries were sequenced on an Illumina NovaSeq 6000 (150 cycles, Illumina).

### Data analysis

Raw single-cell RNA-sequencing data were processed using the BD Rhapsody analysis pipeline (BD Biosciences) on the Seven Bridges platform (https://www.sevenbridges.com) according to the manufacturer’s recommended protocols. The resulting molecule per cell matrix was imported into R version 4.4.3 for subsequent downstream analysis. Initial quality control filtering and normalization were performed using the Seurat R package, version 5.2.1^59^. Low-quality cells were filtered from the dataset, and the top 2,000 most variable genes were selected for downstream analyses. Doublet were identified using the scDblFinder R package, version 1.20.2^60^. A dimensionality reduction was conducted via principal component analysis (PCA) such that the principal components explained up to 90% of the total variance. Cell neighborhoods were identified using Seurat’s FindNeighbors function, and clusters were detected via its FindClusters function. Data visualization was performed using the Uniform Manifold Approximation and Projection (UMAP) technique^61^. Batch effects were corrected using the Harmony R package, version 1.2.3^62^. Cluster-specific marker genes were identified through pairwise differential expression analysis using Seurat’s FindAllMarkers function and the Wilcoxon rank-sum test. Cell type annotations were assigned based on established marker gene databases, including PanglaoDB^63^, the Human Protein Atlas^64^, and the Single Cell Portal^65^. Differential gene expression analysis was performed with statistical thresholds for log2 fold change of greater than 1 and adjusted p-value of less than 0.05. Gene set enrichment analysis was conducted using fgsea version 1.32.4. Cell cycle classification was performed using ccAFv2 version 0.0.0.9^66^. Developmental potential was estimated using CytoTRACE2 version 1.1.0^67^. RNA velocity analysis was conducted using velocyto version 0.17^68^ and scVelo version 0.2.5^69^. Graph-based trajectory inference was performed via Partition-based Graph Abstraction (PAGA)^70^ as implemented in scVelo. All Circos plots were visualized using circlize version 0.4.16^71^

### Flow cytometry

To quantify GFP-expressing cells in organoids generated from *Lgr5-CreERT2-IRES-GFP* mice, the organoids were first dissociated into single cells using TrypLE. The resulting single-cell suspensions were passed through a 40 µm cell strainer and resuspended in PBS containing 0.5% BSA. The cells were then analyzed and sorted using a FACSAria III (BD Biosciences) equipped with a 488 nm laser for GFP detection. GFP-positive and GFP-negative populations were gated relative to unstained controls and analyzed using FlowJo v.10.5.3 (BD Biosciences). The proportion of GFP-positive cells within the total live cell population was quantified.

### Holotomography

Holotomographic images of intestinal organoids were acquired using an HT-X1 system (Tomocube Inc., Daejeon, Korea). Imaging was conducted every six hours for two days under controlled conditions (37□, 5% CO2) using a stage-top incubator to sustain cell viability. After imaging, the organoids were fixed in 4% PFA for volume and crypt quantification.

The organoid volume analysis was conducted using Aivia AI Image Analysis Software v14 (Leica Biosystems), specifically employing the 3D Object Analysis-Meshes recipe. 3D holotomography images were first converted into maximum intensity projection (MIP) images. These MIP images were then imported into Aivia where the 3D Object Analysis-Meshes algorithm segmented the organoids across multiple slices. This process allowed for the quantification of morphological parameters, including volume, surface area, and intensity measurements. The organoid 3D structures were then segmented and stored as ‘Masked Channel’ images using Aivia. To visualize these structures in 2D, a depth-encoded color map was applied to the image stack, followed by maximum intensity projection using the Z-stack Depth Colorcode Fiji plugin. The color map used for each image is indicated by a color bar.

### Statistical analysis

All statistical analyses were performed using GraphPad Prism (GraphPad Software v.8, San Diego, CA, USA). Normality was assessed using the Kolmogorov–Smirnov test. Depending on the distribution of the data, unpaired two-tailed t-tests or nonparametric Mann– Whitney U tests were used for comparisons between two groups. For comparisons among more than two groups, one-way ANOVA with Bonferroni post hoc tests or Kruskal–Wallis tests with Dunn’s or Mann–Whitney U post hoc tests were used. All values are presented as means ± S.E.M. Statistical significance is denoted as follows: \**P* < 0.05, \*\**P* < 0.01, and \*\*\**P* < 0.001.

## Acknowledgements

This work was supported by National Research Foundation of Korea (NRF) grants RS-2023-00208193 and NRF-2022M3A9F3094559 (to Y.T.J.) and RS-2023-00212238 funded by the Korean Government (Ministry of Science and ICT), and by the IITP (Institute of Information & Communications Technology Planning & Evaluation)-ICAN (ICT Challenge and Advanced Network of HRD) grant IITP-2024-RS-2024-00438263 (to J.C.). We appreciate Dr. Peihua Jiang (Monell Chemical Senses Institute) for his generous gift of the stable Noggin-expressing cell line. We appreciate the Institute of Biomedical Science & Food Safety, CJ-Korea University Food Safety Hall (Seoul, Republic of Korea) for providing technical support for this study.

## Author contributions

D. W. S conducted the *in vivo* and organoid experiments. S. S. K. and J. M. C. analyzed the bioinformatic data. H. B. J and M. H. C. conducted the holotomographic image acquisition. M. G. L. and J. H. J. conducted some organoids experiments. J-S. L., S-H. C., and D-H. K. analyzed the data. Y.T.J. supervised the project and wrote the paper. All authors read and approved the manuscript.

