## Supplemental Figures for "Multilayered molecular and cellular mechanisms underlying intestinal epithelial regeneration"

### **(Running title: Multiple mechanisms of intestinal regeneration)**

Dong Woo Seo^1,2^, Seungsoo Kim^2^, Hoibin Jeong^3^, Mi Hyeon Cho^3^, Ming Gyeong Lee^1,2^, Jea Hwa Jang^1,2^, Jun-Seok Lee^1,2^, Sang-Hyun Choi^1^, Dong-Hoon Kim^1,2^, Jungmin Choi^2,*^ and Yong Taek Jeong^1,2,*^

^1^Department of Pharmacology, Korea University College of Medicine, Seoul 02841, Republic of Korea

^2^BK21 Graduate Program, Department of Biomedical Sciences, Korea University College of Medicine, Seoul 02841, Republic of Korea

^3^Metropolitan Seoul Center, Korea Basic Science Institute (KBSI), Seoul 02841, Republic of Korea

*Correspondence

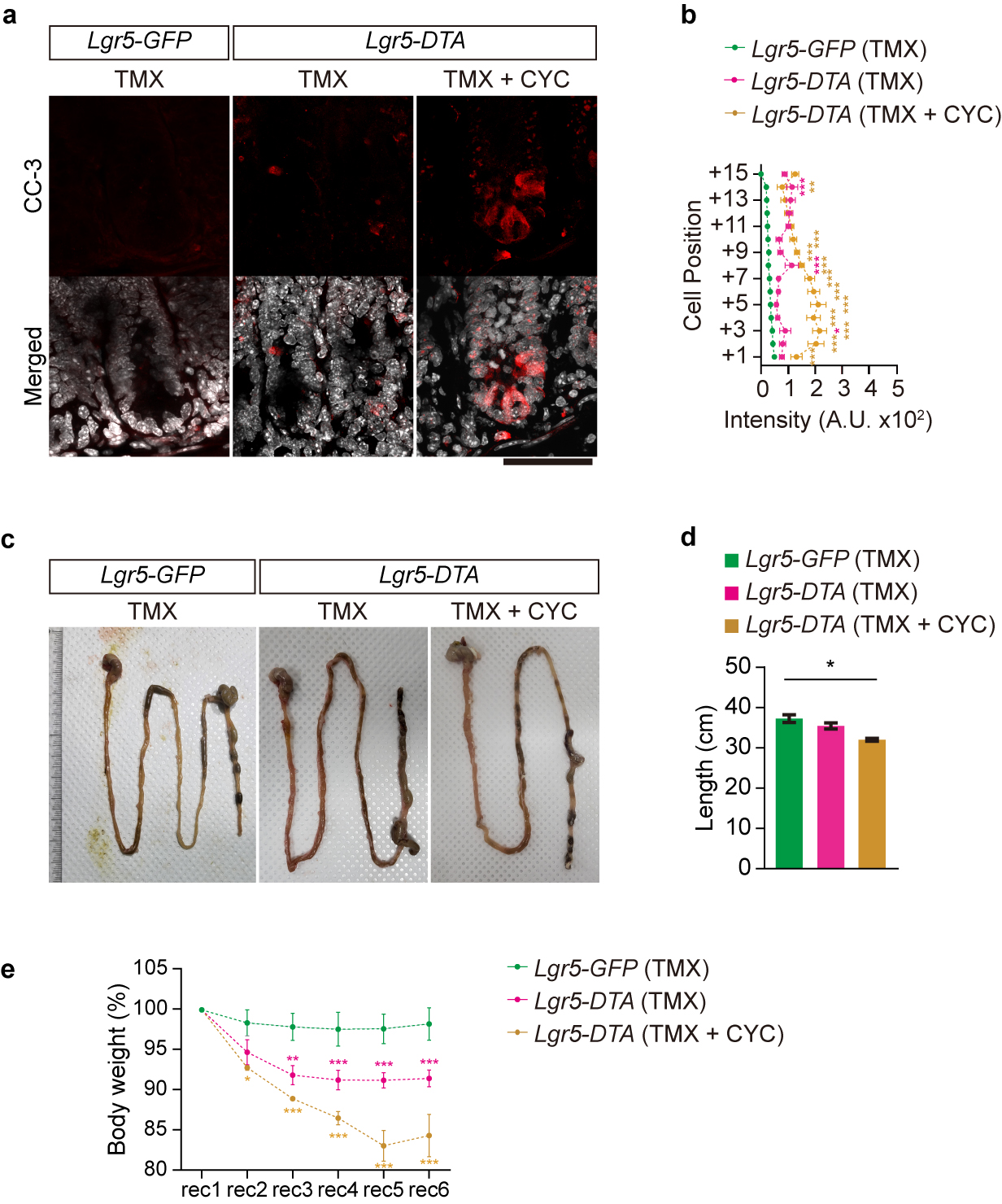
**Extended Data Figure 1 | Hh inhibition attenuates crypt regeneration upon loss of Lgr5+ CBCs. a**, Representative immunofluorescence images showing cleaved caspase-3 (CC-3) staining (red), indicating apoptotic cells in intestinal sections taken on rec 1 from *Lgr5-GFP* and *Lgr5-DTA* mice treated with tamoxifen (TMX) or TMX and cyclopamine (CYC). Scale bars: 50 µm. **b**, Quantification of CC-3 fluorescence intensity across cell positions in the crypt on rec 1 (*n* = 5). **c**, Representative images showing the morphology of the gastrointestinal tract of *Lgr5-GFP* and *Lgr5-DTA* mice treated with TMX or TMX+CYC (*n* = 4). **d**, Average intestinal length on rec 6 of the treatment period (*n* = 4). e, Changes in the body weight of *Lgr5-GFP* and *Lgr5-DTA* mice after treatment (from rec 1 to rec 6) with TMX or TMX+CYC (*n* = 5). All data are presented as means ± S.E.M. Two-way ANOVA was used to assess differences among groups with Tukey’s post hoc test for multiple comparisons in (**b**, **e**) or Mann-Whitney (two-tailed) U-tests in (**d**). **P* < 0.05, ***P* < 0.01, and ****P* < 0.001.

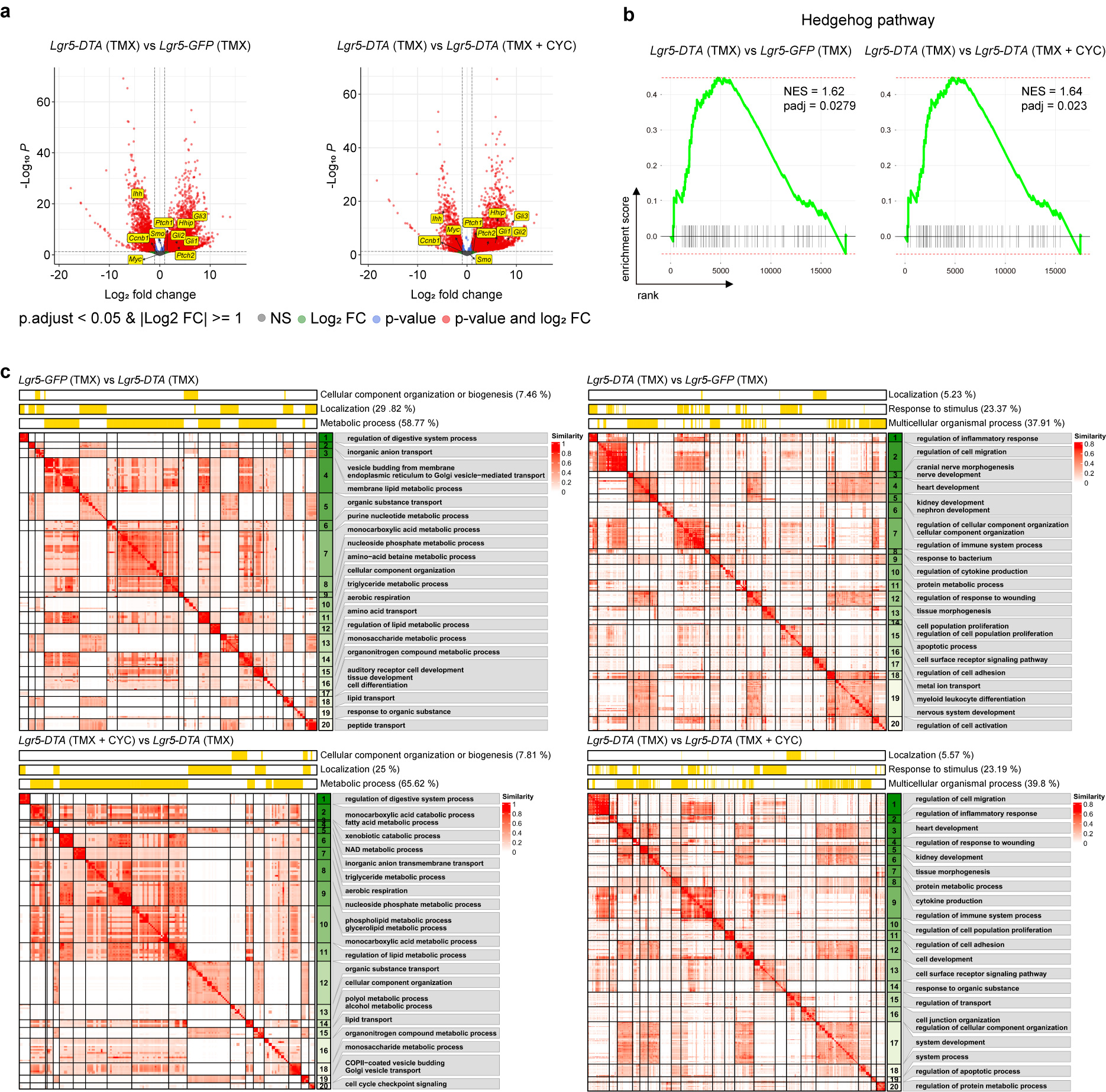
**Extended Data Figure 2 | Hh inhibition attenuates regenerative transcriptomic responses upon loss of Lgr5+ CBCs. a**, Volcano plots showing DEGs in the comparison of *Lgr5-DTA* (TMX) and *Lgr5-GFP* (TMX) (left) or *Lgr5-DTA* (TMX) and *Lgr5-DTA* (TMX+CYC) (right). Genes with an adjusted p-value < 0.05 and an absolute log₂ fold change (log₂FC) ≥ 1 are highlighted in red. The genes labeled with a yellow box are those that are up-regulated by the Hh signaling pathway. **b**, GSEA plot showing enrichment of the Hh signaling pathway in *Lgr5-DTA* (TMX) compared to other groups. **c**, Heatmap displaying GSEA results from differential expression analysis across four pairwise comparisons. For each comparison, significantly enriched Gene Ontology Biological Process (GOBP) terms (FDR < 0.05) were clustered based on semantic similarity using in-house scripts. Term similarity is represented by the intensity of red. The representative term for each cluster is shown in the gray box to the right. Clusters are ranked from 1 to 20 according to descending average normalized enrichment score (NES), with green intensity reflecting average NES values. *Lgr5-GFP* (TMX) vs *Lgr5-DTA* (TMX) (top left), *Lgr5-DTA* (TMX) vs *Lgr5-GFP* (TMX) (top right), *Lgr5-DTA* (TMX+CYC) vs *Lgr5-DTA* (TMX) (bottom left) and *Lgr5-DTA* (TMX) vs *Lgr5-DTA* (TMX+CYC) (bottom right). The annotation panel above each heatmap summarizes broader GOBP categories, and the percentages indicate the proportion of child terms assigned to each category.

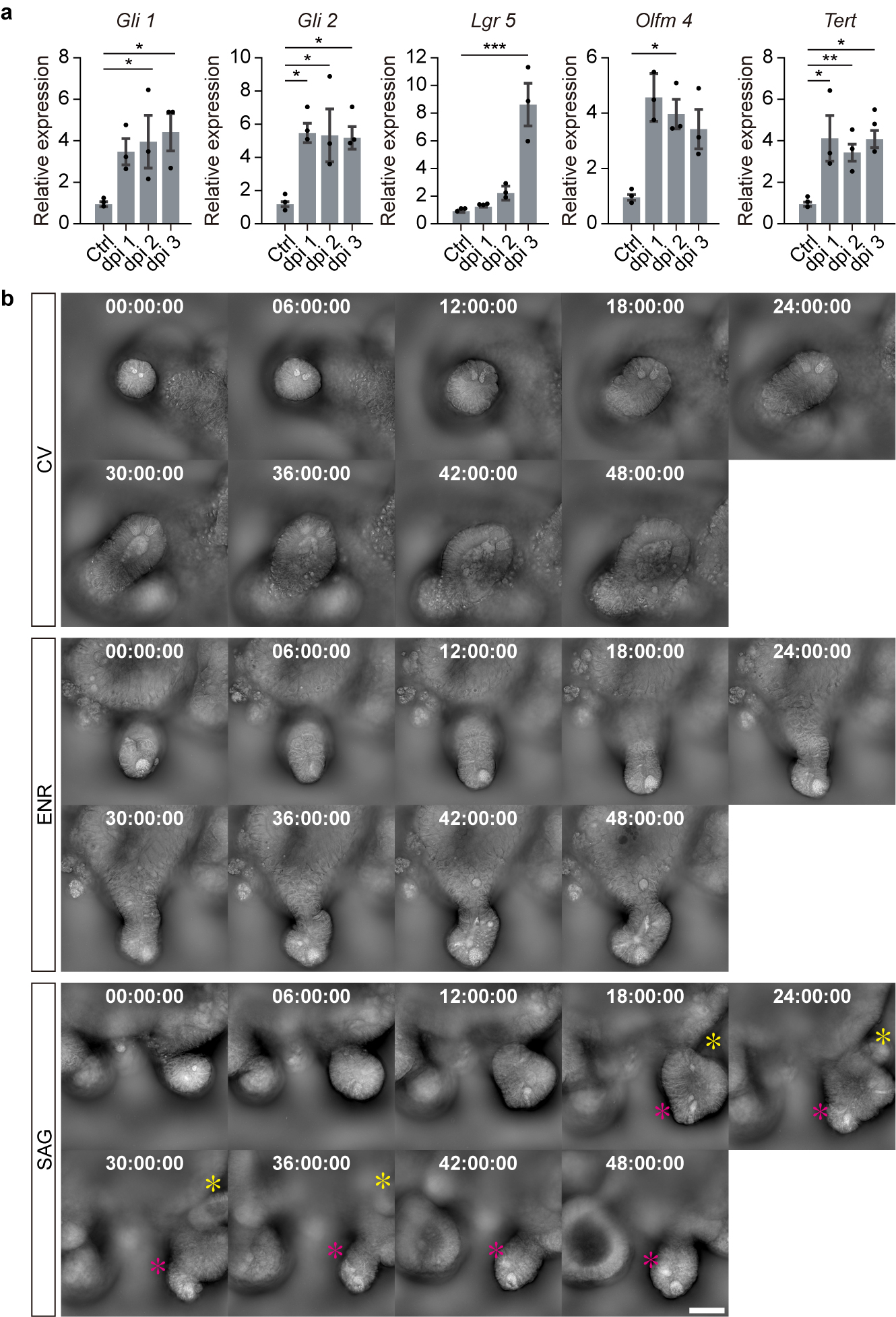
**Extended Data Figure 3 | Hh activation induces up-regulation of stem cell signature genes and the emergence of de novo crypts. a**, qRT-PCR analysis of the relative mRNA expression levels of *Gli1*, *Gli2*, *Lgr5*, *Olfm4*, and *Tert*. RNA was isolated from the crypts of control and SAG-treated groups on dpi 1, dpi 2, and dpi 3. Data are presented as means ± S.E.M. (*n* = 3 mice per group). One-way ANOVA with Tukey’s post hoc test. **P* < 0.05, ***P* < 0.01, and ****P* < 0.001. **b**, Time-lapse images were captured every 6 hours for a total of 48 hours to observe crypt changes in organoids treated with CV, ENR, or SAG. Yellow stars indicate de novo crypt formation, while pink stars indicate pre-existing crypts. Scale bars: 50 µm.

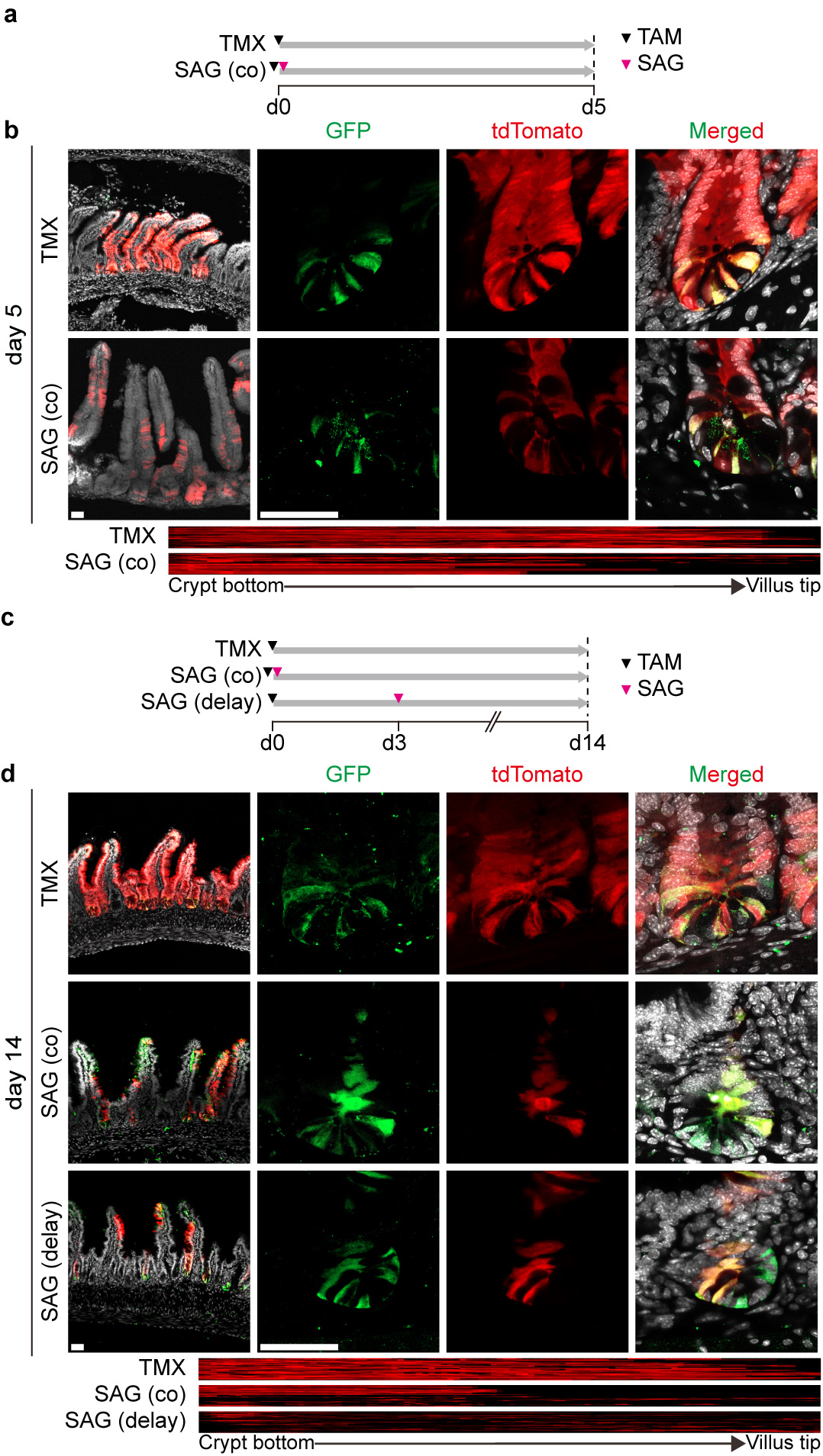
**Extended Data Figure 4 | Delayed SAG treatment enhances the emergence of de novo Lgr5⁺ CBCs from non-Lgr5⁺ clones. a**, Timeline of the experimental design for (**b**). **b**, Top: Representative images of crypt-villus compartments on day 5 after treatment with TMX alone or TMX with SAG. Bottom: Heatmap showing the normalized distribution of tdTomato from the crypt bottom to the villus tip (TMX: *n* = 35 crypt-villus units, SAG (co): *n* = 30, from 5 mice each). **c**, Timeline of the experimental design for (**d**). **d**, Top: Representative images of the crypt–villus compartment on day 14 across three groups: TMX, SAG (co) (co-treated with TMX), and SAG (delay) (SAG injected 3 days after TMX injection). Bottom: Heatmap showing the normalized distribution of tdTomato from the crypt bottom to the villus tip (bottom) (TMX: *n* = 23, SAG (co): *n* = 32, and SAG (delay) *n* = 43, from 5 mice each). Scale bars: 50 µm (**b**, **d**).

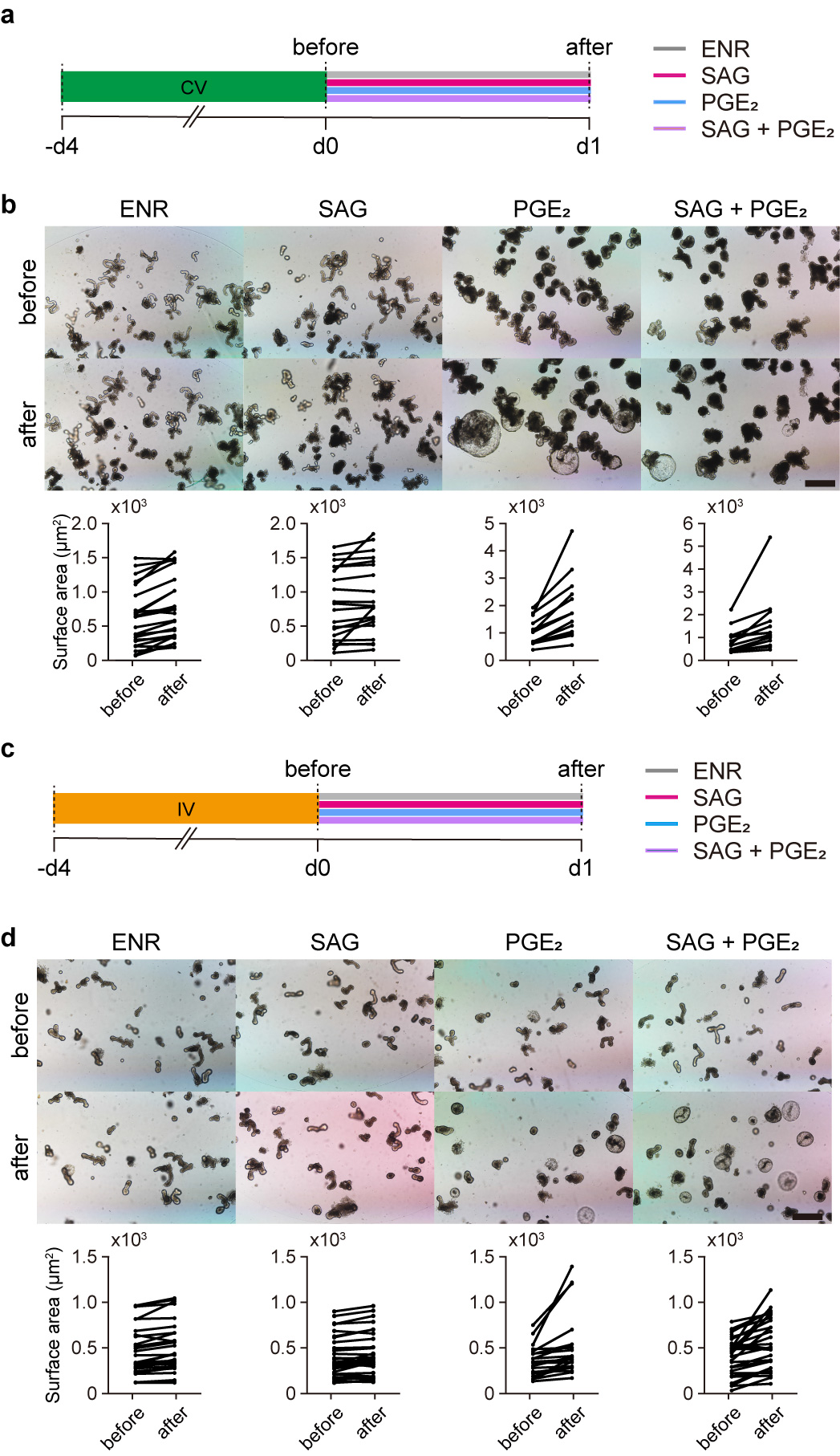
**Extended Data Figure 5 | SAG and PGE_2_ induce distinct regenerative responses in crypts and villi, respectively. a**, Experimental timeline for CV-primed organoids. **b**, Representative images and surface area quantification before and 24 hours after treatment with ENR, SAG, PGE_2_, or SAG + PGE_2_ in organoids primed with CV (ENR: *n* = 23, SAG: *n* = 21, PGE_2_: *n* = 18, SAG + PGE_2_: *n* = 19). **c**, Experimental timeline for IV-primed organoids. **d**, Representative images and surface area quantification before and after treatment with ENR, SAG, PGE_2_, or SAG + PGE_2_ in organoids primed with IV (ENR: *n* = 29, SAG: *n* = 32, PGE_2_: *n* = 22, SAG + PGE_2_: *n* = 33). Scale bars: 200 µm (**b**, **d**).

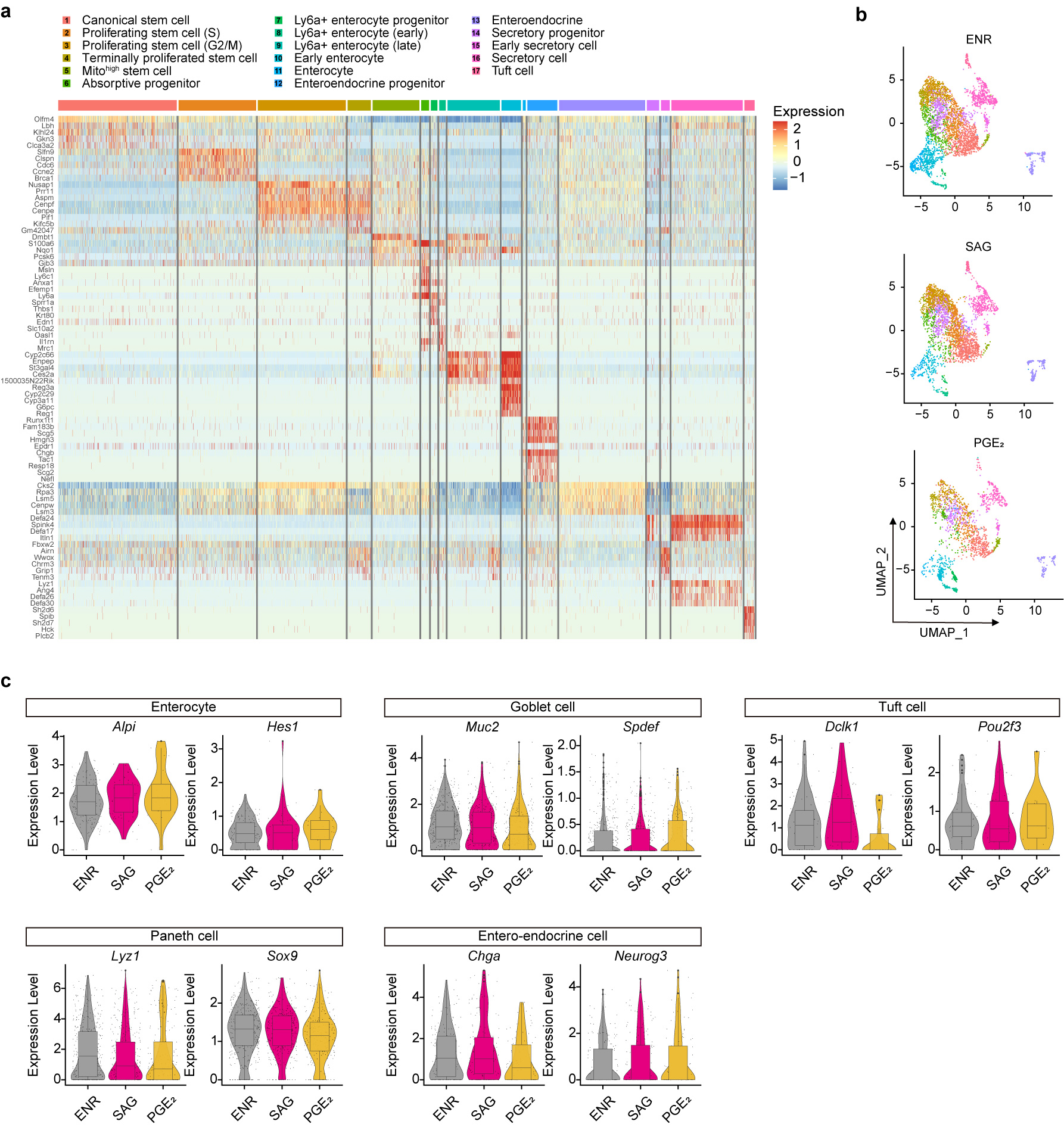
**Extended Data Figure 6 | Annotation and validation of intestinal organoid scRNA-seq. a**, Heatmap showing the top five marker genes for each cell cluster, revealing distinct transcriptional signatures across differentiation-defined populations in intestinal organoids. Cell type identities are color-coded along the top bar. **b**, UMAP plots visualizing cell clusters from scRNA-seq datasets of organoids cultured in ENR-, SAG-, or PGE_2_-conditions. **c**, Violin plots showing the expression levels of key marker genes for enterocytes (*Alpi*, *Hes1*), goblet cells (*Muc2*, *Spdef*), Paneth cells (*Lyz1*, *Sox9*), tuft cells (*Dclk1*, *Pou2f3*), and enteroendocrine cells (*Chga*, *Neurog3*) under each culture condition.

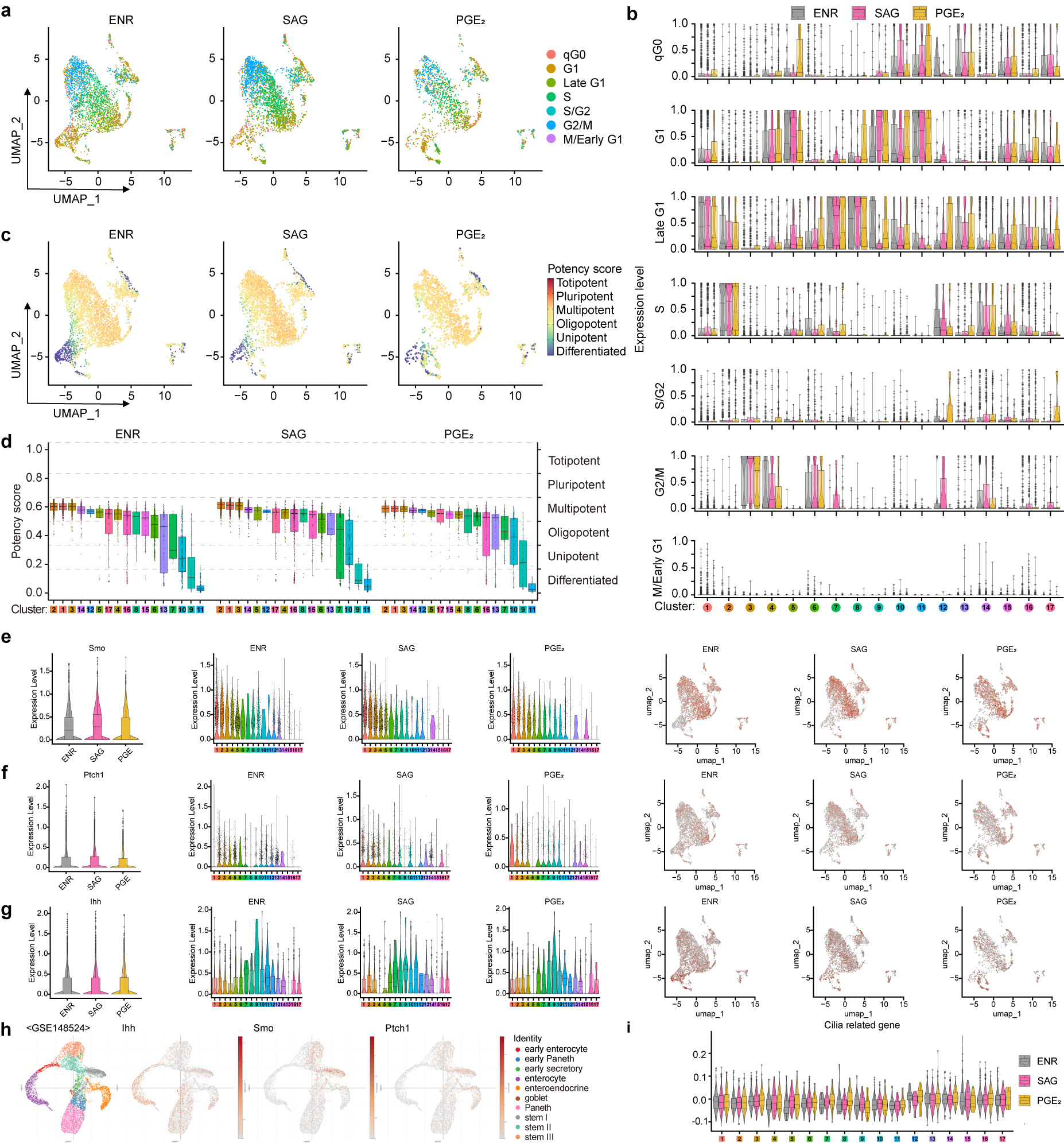
**Extended Data Figure 7 | Hh activation promotes cell cycle progression and enhances differentiation potential, as well as increasing Hh pathway gene expression. a**, UMAP plots showing the distribution of cell cycle phases, as inferred by the ccAFv2 algorithm, across ENR-, SAG-, and PGE_2_-treated organoids. **b**, Violin plots displaying the distribution of annotated cell types for each cell cycle phase under each condition. **c**, UMAP plots colored by predicted potency states, ranging from totipotent to differentiated, as determined by the CytoTRACE2 package. **d**, Box plots of CytoTRACE2-inferred potency scores across annotated cell clusters, arranged in decreasing order of their CytoTRACE2 scores. **e**–**g**, Expression of Hh pathway components. Violin plots (left) show the expression levels of *Smo* (**e**), *Ptch1* (**f**), and *Ihh* (**g**) across all clusters for each condition. Violin plots (middle) display the expression of each gene across individual cell clusters for each condition. UMAP feature plots (right) showing the distribution of gene expression within the organoid landscape for each treatment. **h**, UMAP plots showing the expression of Smo, Ptch1, and Ihh in intestinal organoids from a public dataset (GSE148528). **i**, Violin plots of module scores for cilia-related genes across annotated clusters in ENR-, SAG-, or PGE_2_-treated organoids.

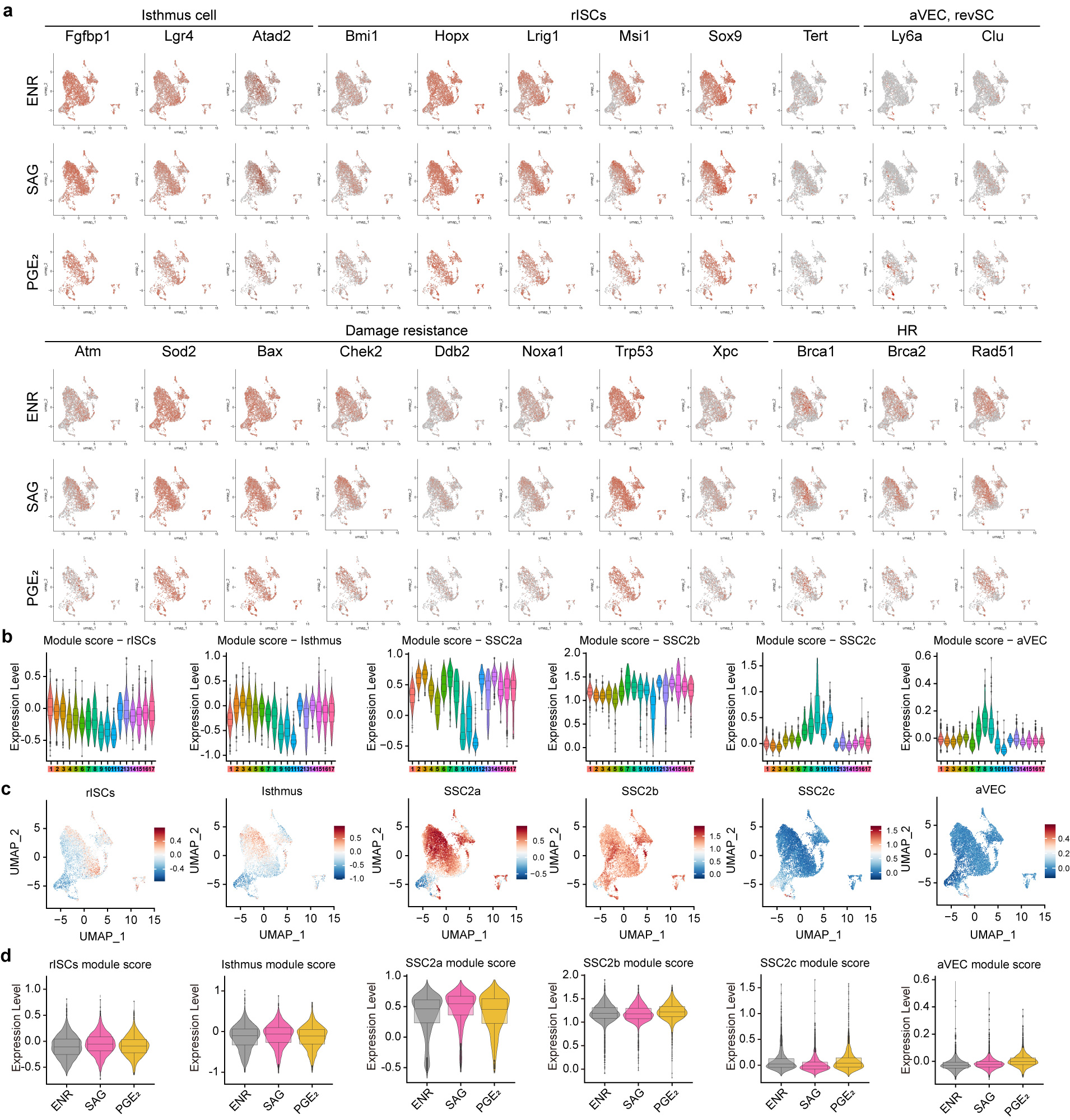
**Extended Data Figure 8 | SAG and PGE_2_ induce distinct regenerative gene programs. a**, UMAP plots showing the expression of markers for isthmus cells, reserve intestinal stem cells (rISCs), atrophy-induced villus epithelial cells (aVECs), revival stem cells (revSCs), and genes related to damage resistance and homologous recombination (HR) across ENR-, SAG-, and PGE_2_-treated organoids. **b**, Module scores for rISCs, isthmus cells, revSCs (comprising SSC2a, SSC2b, and SSC2c), and aVECs across annotated clusters. **c**, UMAP plots showing module score distributions for the indicated regenerative cellular states. **d**, Violin plots showing module score distributions for the indicated regenerative cellular states across the ENR, SAG, and PGE_2_ treatment conditions.

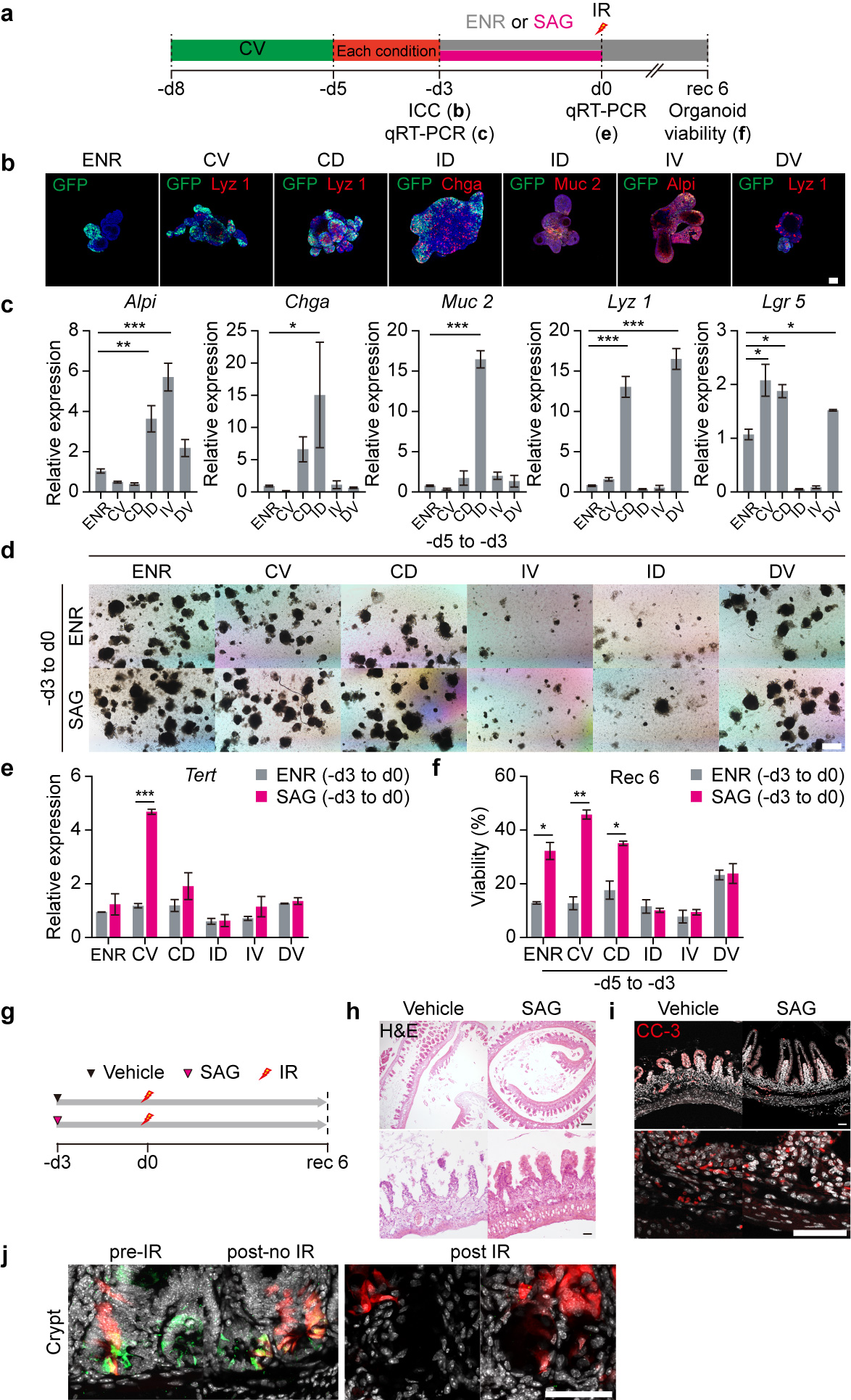
**Extended Data Figure 9 | SAG pre-treatment improves post-irradiation regeneration and viability in vitro and in vivo. a**, Experimental timeline showing in vitro SAG treatment, irradiation (IR), and analysis on day 6 post-IR (rec 6). **b**, Representative immunofluorescence images showing GFP (green) and Lyz1, Chga, Muc2, or Alpi (red) expression under the indicated conditions. C, CHIR-99021; I, IWP-2; D, DAPT; V, Valproic Acid, **c**, qRT-PCR analysis of *Alpi*, *Chga*, *Muc2*, *Lyz1*, and *Lgr5* mRNA levels (*n* = 3). **d**, Bright-field images of irradiated organoids (day 6) post-ENR or SAG treatment. **e**, qRT-PCR quantification of Tert mRNA expression (*n* = 3). **f**, Viability of irradiated organoids post-treatment (*n* = 3). **g**, In vivo timeline of SAG treatment, IR, and tissue harvest on day 6. **h**, Hematoxylin and eosin (H&E) staining of intestinal sections post-IR in the vehicle and SAG groups (×4 and ×20). **i**, Cleaved caspase-3 (CC-3) immunofluorescence showing apoptotic cells (×20 and ×63). **j**, High-magnification crypt images (from **Fig. 5j**). All data are presented as means ± S.E.M. One-way ANOVA followed by Tukey’s post hoc test for (**c**, **e**), and two-way ANOVA with multiple comparisons for (**f**). **P* < 0.05, ***P* < 0.01, and ****P* < 0.001. Scale bars: 50 µm (**b**, **h**–**j**), 200 µm (**d**).

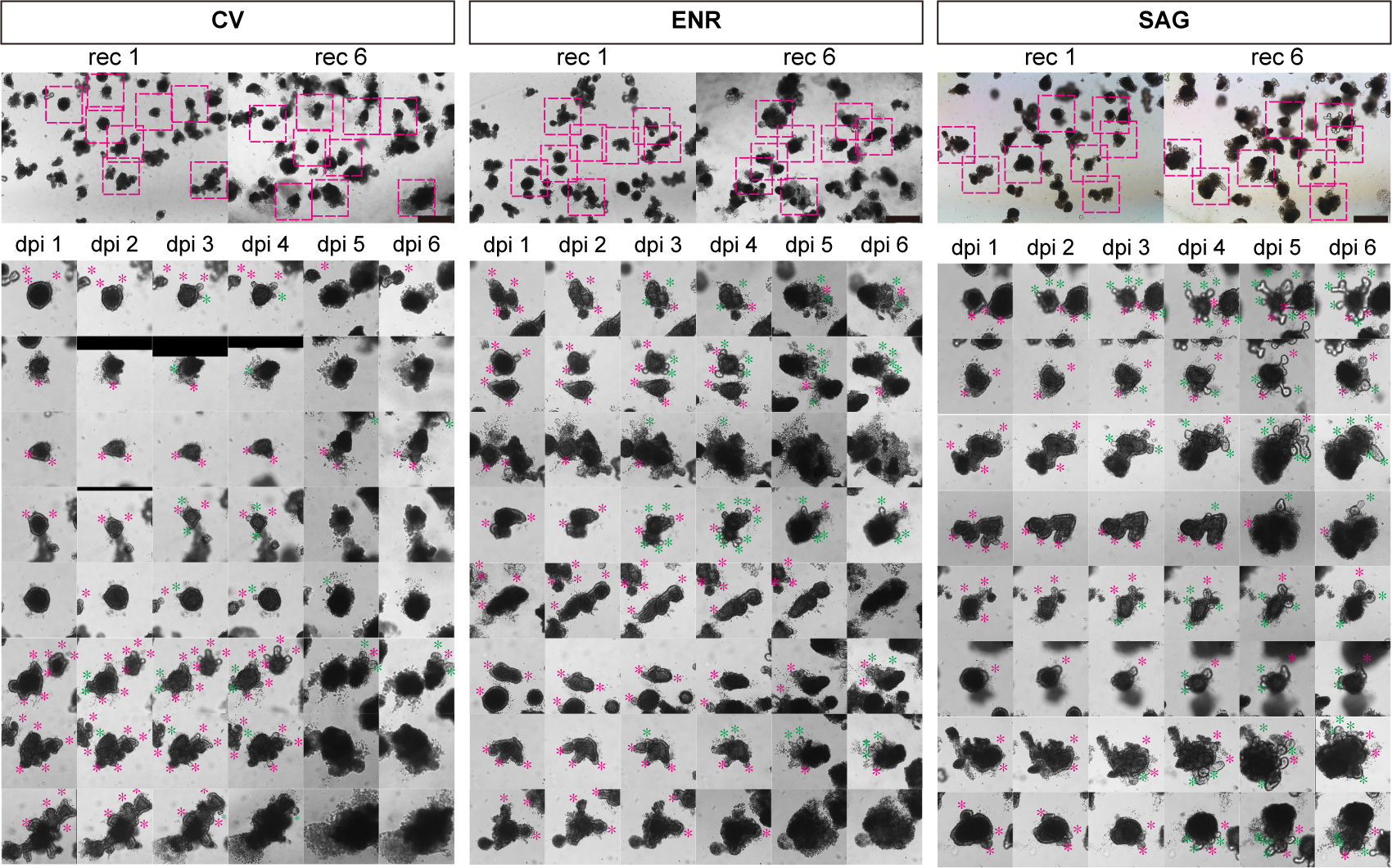
**Extended Data Figure 10 | SAG confers damage resistance on existing crypts and promotes de novo crypt regeneration.** Organoids cultured with CV, ENR, or SAG were irradiated, and their changes were tracked from post-irradiation day 1 to 6. Pink stars indicate pre-existing crypts, and green stars indicate de novo crypts. Scale bars: 20 µm.
